# IgLON5 autoimmune antibodies activate Tau via neuronal hyperactivity

**DOI:** 10.1101/2024.03.10.584272

**Authors:** Bilge Askin, Cagla Kilic, César Cordero Gómez, Sophie Lan-Linh Duong, Alvaro Domingues-Baquero, Alexander Goihl, Karsten Nalbach, Joana Petushi, Pia Grundschöttel, Jessica Wagner, Valentine Thomas, Janne Lamberty, Emily Withers, Hanna Huber, Sabrina Huebschmann, Ekaterina Semenova, Paul Turko, Andrew G. Newman, Lisa Diez, Marc Beyer, Elena De Domenico, Peter Körtvelyessy, Dirk Reinhold, Anja Schneider, Jonas J. Neher, Thomas Ulas, Stefan F. Lichtenthaler, Benjamin R. Rost, Dietmar Schmitz, Harald Prüss, Susanne Wegmann

## Abstract

Anti-IgLON5 disease is an autoimmune disease, in which autoantibodies (AABs) against the neuronal cell surface protein IgLON5 lead to profound brain dysfunction and Tau pathology. How α-IgLON5 AABs cause neuronal Tau protein pathology and neurodegeneration remains unclear. We find that patient-derived α-IgLON5 AABs cluster IgLON5 proteins with other cell surface proteins, leading to neuronal hyperactivity that triggers pathological Tau missorting and phosphorylation, typically observed early in Tau-related neurodegenerative diseases. In wildtype mice, α-IgLON5 AABs induce hippocampal Tau phosphorylation and neuroinflammatory responses. Our findings establish a causal link between the α-IgLON5 AABs and Tau pathology in anti-IgLON5 disease patients, and highlight the role of neuronal hyperactivity as a disease-overarching driver of Tau pathology and provide a potential target for therapeutic intervention.

**Teaser:** α-IgLON5 autoantibodies induce clustering of neuronal cell surface proteins, leading to acute neuronal hyperactivity and Tau missorting.

## Introduction

In autoimmune encephalitis disorders, autoimmune antibodies (autoantibodies, AABs) directed against neuronal or glial epitopes in the brain can cause or promote neuroinflammation and neurodegeneration (*1*, *2*). Anti-IgLON5 disease is a rare but severe neurological disorder characterized by the presence of AABs targeting the neuronal cell surface protein IgLON5 (*3*). Individuals diagnosed with anti-IgLON5 disease demonstrate diverse clinical manifestations, commonly experiencing sleep behavior abnormalities, movement disorders, memory deficits, and seizures (*4–6*). Brains of anti-IgLON5 patients show intraneuronal accumulations of phosphorylated Tau protein in several regions, including brain stem, hypothalamus, cerebellum, hippocampus, and basal ganglia, as well as in the spinal cord (*7*, *8*). Similar neuronal Tau accumulation and aggregation are well-known as pathological hallmarks in primary and secondary tauopathies, including frontotemporal dementia (FTD) variants and Alzheimer’s disease (AD) (*9*, *10*). However, anti-IgLON5 disease differs from other tauopathies because of its unique autoimmune context, anatomical distribution of Tau pathology, and set of clinical manifestations (*3*). The occurrence of Tau pathology in anti-IgLON5 disease poses the fundamental questions of how AABs against a cell surface protein can trigger pathophysiological intracellular Tau changes.

IgLON5 is primarily expressed in neurons in the brain stem, medulla oblongata, thalamus, cerebral cortex and cerebellum, and to a lesser degree in other brain areas like the hippocampal formation and amygdala (*11*). It belongs to the IgLON protein family, a group of immunoglobulin (Ig) domain cell adhesion molecules comprising five members (OPCML (=IgLON1), NTM (=IgLON2), LSAMP (=IgLON3), NEGR1 (=IgLON4), and IgLON5)) (*12*). IgLON5 is a GPI-anchored surface protein with three Ig-domains (Ig1, Ig2, and Ig3) that extend into the extracellular space and mediate homo- and heterodimer formation with other IgLON family members on the cell surface (*13–15*). The physiological function of IgLON5 is not fully understood, however, other IgLON family members have been implicated in neuronal cell adhesion, neurite growth, neural circuit and synapse formation (*16–18*). In neuronal cultures, binding of AABs from anti-IgLON5 patient serum to surface IgLON5 was shown to induce protein-AAB complex internalization (*19*), and long-term treatment (up to 4 weeks) induced common neurodegenerative phenotypes like cytoskeleton disruption (*20*), synapse loss and cell death (*21*).

Here, using purified α-IgLON5 AAB from anti-IgLON5 disease patient plasma, we show that α-IgLON5 AABs promote pro-pathological Tau changes and neurotoxicity by inducing acute neuronal hyperactivity. The present findings establish an important novel molecular and cellular mechanism underlying Tau pathology in anti-IgLON5 disease, which in turn provides a foundation for the development of treatment strategies aimed at preventing Tau changes in autoimmune encephalitis patients.

## Results

### α-IgLON5 AABs isolated from patient plasma

To investigate the effects of α-IgLON5 AABs on neurons and Tau, we isolated polyclonal pools of IgLON5-specific AABs (α-IgLON5 AABs) from the plasma of four anti-IgLON5 disease patients through affinity chromatography with immobilized recombinant human IgLON5-Fc chimera (Fig. 1A). The purified α-IgLON5 AABs had a purity of 78±6% (mean±SEM; signal intensity of heavy (∼50 kDa) and light chain (∼25 kDa) relative to total signal in SDS-PAGE; Fig. 1B). Immunoblotting confirmed the binding of the isolated α-IgLON5 AAB pool to recombinant IgLON5 protein but not to LGI1, another neuronal surface protein (fig. S1A), as well as the absence of residual IgLON5 protein in the AAB preparations (fig. S1B). The IgG subclass composition of isolated α-IgLON5 AAB pools (by ELISA) appeared to be similar for all patient α-IgLON5 AAB preparations (fig. S1C), with 37-50% being IgG_1_, 37-52% IgG_2_, 2-3% IgG_3_, and 8-13% IgG_4_. IgG pools derived from healthy subject serum (pCtrl) or commercially available human serum (IgG_pool_) showed similar proportion of IgG subclasses as well. Dot blot analysis of IgG subclasses confirmed the ELISA data for α-IgLON5#1 and pCtrl (fig. S1D). Additionally, we found that the IgM levels were comparable across samples, with a slight increase observed in α-IgLON5#4 and pCtrl (fig. S1E). pCtrl antibodies were subsequently used throughout the manuscript as control IgG pool.

**Fig. 1.**
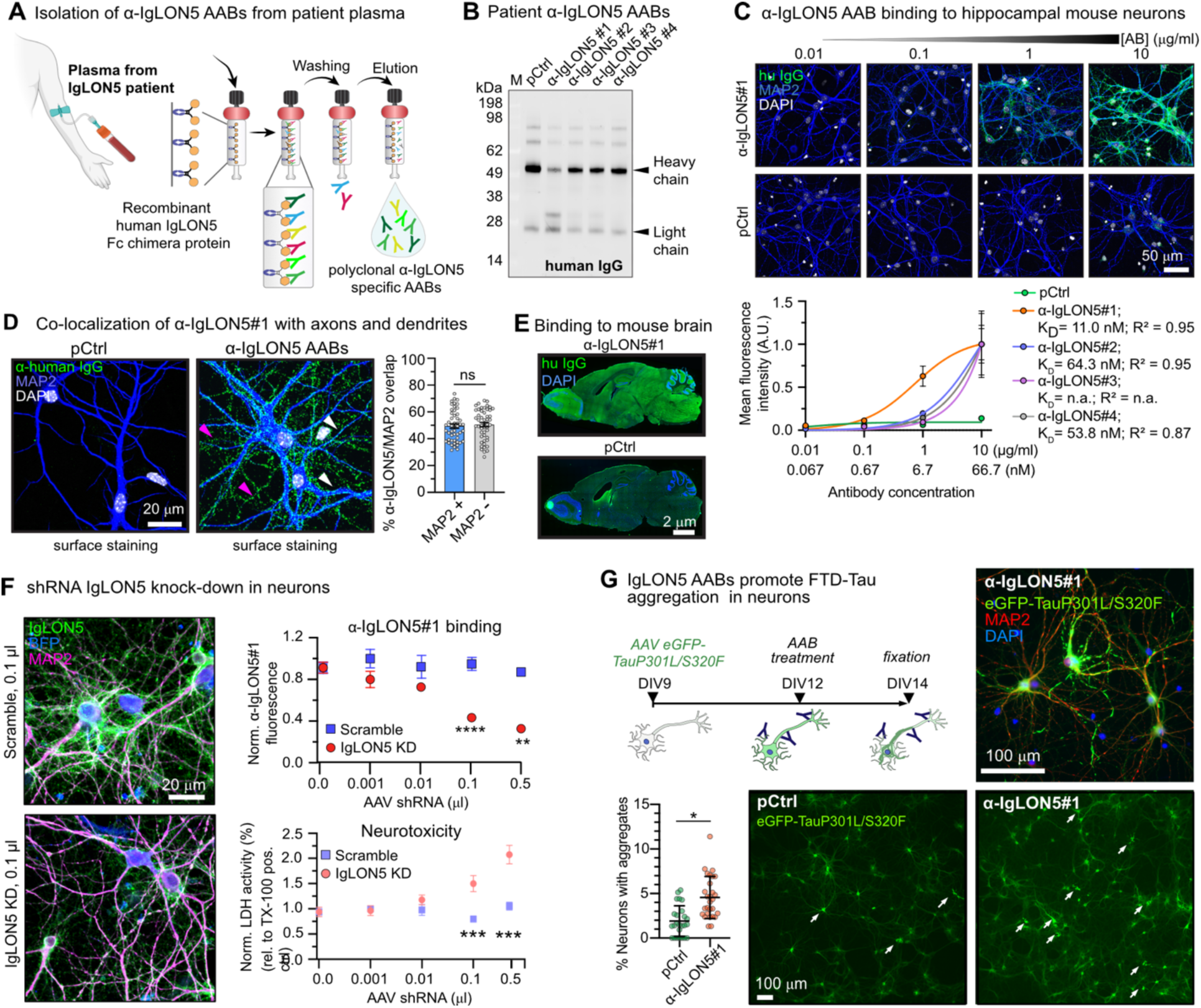
α-IgLON5 AABs from patient plasma bind neuronal IgLON5. (**A**) Principle of α-IgLON5 AAB isolation from clinically verified anti-IgLON5 patient plasma using affinity purification on immobilized recombinant human IgLON5 Fc-chimera. (**B**) Western blot of patient α-IgLON5 AABs (IgLON5#1-4) and pCtrl probed with α-human IgG shows the presence of IgG heavy and light chains. (**C**) Dose-dependent binding of α-IgLON5 AABs and pCtrl to PFA-fixed primary hippocampal mouse neurons. K_D_’s were calculated via non-linear fitting. (**D**) Binding of α-IgLON5#1 to dendrites (MAP2+; white arrows) and axons (MAP2-; pink arrows). N = 3 experiments with 3-5 repeats per condition. Data points represent individual images analyzed. Student’s t-test. (**E**) Immunoreactivity of α-IgLON5#1 and pCtrl in fresh-frozen, sagittal brain sections of an adult wildtype mouse. (**F**) shRNA-mediated IgLON5 knockdown in primary neurons. Quantification of α-IgLON5#1 binding to neurons transduced with serial dilutions of AAVs encoding anti-mIgLON5 or scrambled shRNA. mIgLON5 shRNA AAV dose-dependent neurotoxicity (LDH assay) occurs ≧0.1 μl shRNA AAVs. AAB signal was measured from dendrites (MAP2+ area). N = 3 experiments with 3-5 replicates. One-way ANOVA with Tukey post-rest. (**G**) Experimental setup, representative images and quantification of NFT-like Tau aggregation in neurons expressing eGFP-TauP301L/S320F for 5 days, then treated with 1 μg/ml α-IgLON5#1 or pCtrl for 2 days. White arrows indicate Tau NFTs neurons. N = 3 experimental replicates. Data points are individual images. Student’s t-test. All panels: Data shown as mean ± SEM. Scale bars as indicated.

### Patient-derived α-IgLON5 AABs bind neuronal cell surface IgLON5

To examine the surface antigen affinity of α-IgLON5 AABs, we measured their binding to the neuronal cell surface. Staining of PFA-fixed, unpermeabilized primary hippocampal mouse neurons (DIV12) with α-IgLON5 AABs revealed different binding efficiencies, measured based on anti-human secondary antibody immunofluorescence: α-IgLON5#1 (K_D_ = 11 nM) showed the strongest and α-IgLON5#3 the weakest binding (Fig. 1C; fig. S1F). Further, we found that α-IgLON5 AABs (exemplified for α-IgLON5#1), bound similarly to cell bodies, dendrites (MAP2+) and axons (MAP2-) (Fig. 1D), and did not show a preference for pre-(Synapsin-1) or postsynaptic (PSD95) areas (fig. S1G). We could also confirm reactivity of α-IgLON5#1 in mouse brain sections (Fig. 1E) and on the surface of cultured human neurons (fig. S1H). To confirm the specificity of α-IgLON5 AABs for IgLON5 protein, we confirmed dose-dependent binding of α-IgLON5#1 to HEK cells recombinantly expressing human IgLON5 protein (fig. S1I). Furthermore, in mouse neurons with shRNA-mediated IgLON5 knock-down (IgLON5 KD), the binding of α-IgLON5#1 decreased in a shRNA-dose dependent manner (Fig. 1F). IgLON5 KD in mouse neuroblastoma cells (Neuro2a) confirmed the efficiency of the IgLON5 shRNA approach (fig. S1J). Notably, the reduction of neuronal surface IgLON5 by ≥30% (at ≥0.1 μl AAV shRNA) appeared to be neurotoxic (Fig. 1F), a finding consistent with previous reports (*19*). Finally, to determine which part of IgLON5 protein was bound by α-IgLON5 AABs, we titrated α-IgLON5#1 to HEK cells recombinantly expressing human full-length IgLON5 or (combinations of) its individual GPI-anchored Ig-domains (Ig1/2/3) (fig. S1K). These experiments showed binding of the polyclonal α-IgLON5#1 pool to all three Ig domains. Notably, previous studies suggested binding of non-purified IgLON5 AABs in patient serum mostly to Ig2 (*19*).

### α-IgLON5 AABs induce Tau missorting

The majority of IgLON5 patients develops neuronal Tau accumulation and aggregation in one or more brain regions (*3*, *7*, *22*). To test whether α-IgLON5 AABs would be able to promote Tau aggregation, we applied α-IgLON5#1 on neurons that recombinantly expressed (AAV mediated) human Tau FTD-mutants with increased aggregation potential. In neurons expressing TauP301L or TauΔK280, α-IgLON5#1 treatment was unable to set-off Tau aggregation (fig. S2A). However, in neurons expressing the spontaneously aggregating FTD-double mutant TauP301L/S320F (*23*, *24*), α-IgLON5#1 treatment induced a significant increase in neurofibrillary tangle-like Tau aggregate formation compared to pCtrl treated neurons (α-IgLON5#1: tangles in 5% of transduced neurons; pCtrl: tangles in 2% neurons; Fig. 1G). α-IgLON5 AABs therefore promoted spontaneous FTD-Tau aggregation.

Missorting of phosphorylated Tau from the axon into the soma and dendrites– is one of the earliest signs of Tau “activation” in the context of neuronal stress and neurodegenerative diseases like AD (*25*). Prolonged Tau missorting is thought to precede pathological Tau oligomerization and aggregation in AD and tauopathies (*26*). To assess whether α-IgLON5 AABs were able to induce *a priori* Tau missorting, we treated hippocampal neurons with the different α-IgLON5 AABs for 2 days and assessed the levels of endogenous Tau in neuronal somata by immunofluorescence. Treatment with α-IgLON5#1 for 2 days induced somatic Tau accumulation in a dose dependent manner (Fig. 2A, B; fig. S2B) and mild neurotoxicity over time (fig. S2C). Notably, overall Tau protein levels, as determined by pan-neuronal immunofluorescence and Western Blot of neuronal lysates, were unchanged by α-IgLON5 treatment, indicating no transcriptional or translational upregulation of Tau (fig. S2D, E). High α-IgLON5#1 concentration (50 μg/ml) induced fragmentation of neuronal processes already after 2 days (Fig. 2A), indicating specific or unspecific neurotoxicity. α-IgLON5#1 therefore exerted a time- and dose-dependent toxicity. Similarly, prolonged treatment of cultured neurons with the total IgG fraction from α-IgLON5 patient serum - containing high doses of unpurified antibodies (e.g., 10-50 μg/ml AABs for 5-21 days) - was reported to induce neurotoxicity (*19*, *21*).

**Fig. 2.**
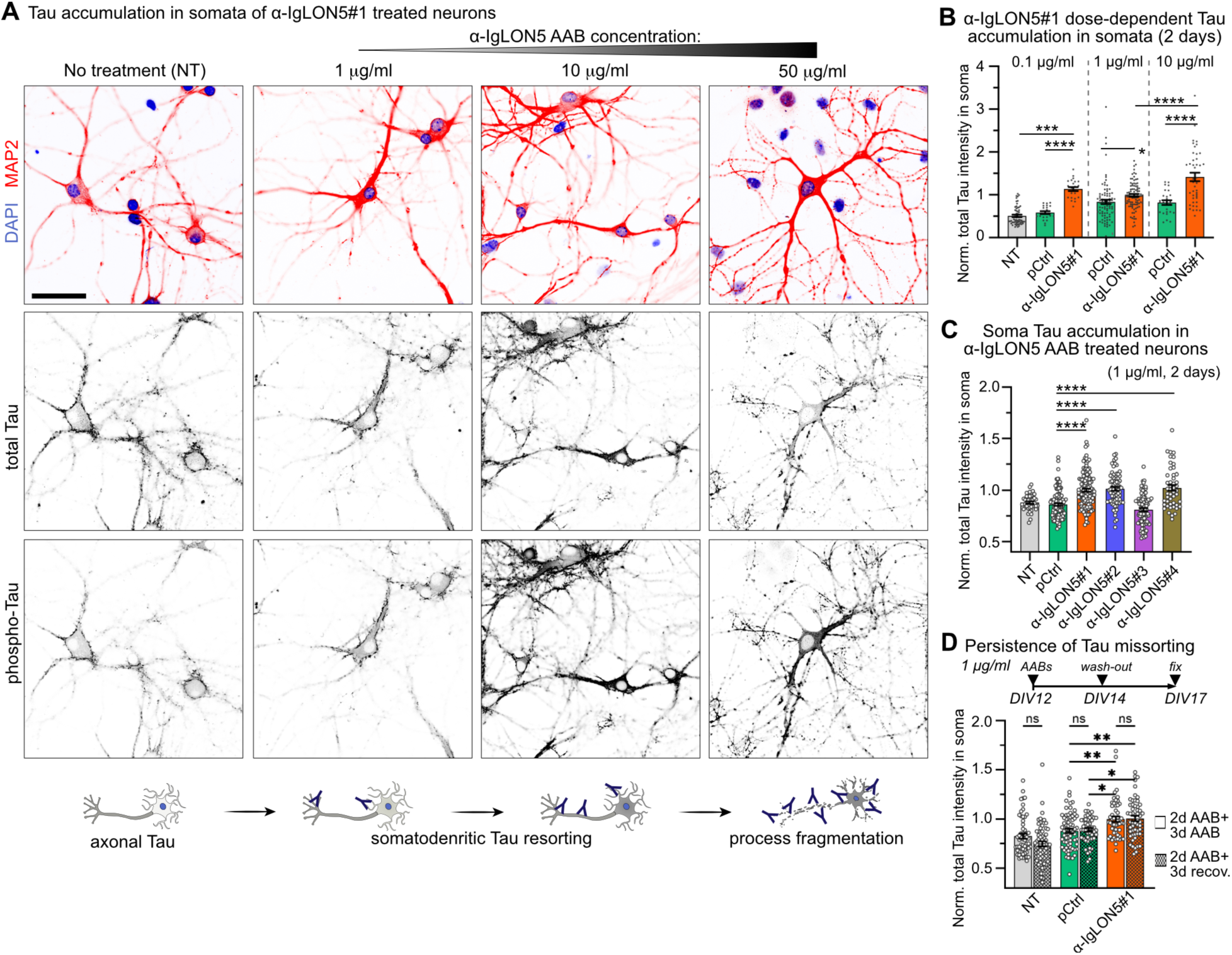
α-IgLON5 AABs promote somatodendritic Tau missorting. (**A**) Hippocampal neurons treated with increasing α-IgLON5#1 concentration (1, 10, and 50 μg/ml) for 2 days, immunolabeled for MAP2, total Tau, and phosphorylated Tau (pS199/pT205/pT231/pS396). Untreated neurons (NT) are shown for comparison. Treatment with 1 and 10 μg/ml α-IgLON5#1 showed increased total Tau and phospho-Tau in neuronal cell bodies. At 50 μg/ml, fragmentation of neuronal processes indicates neurotoxicity and neurons showed strong phospho-Tau reactivity in cell bodies and elevated signal in the nucleus, whereas total Tau decreased in cell bodies. Scale bar is 50 μm. (**B**) Quantification of total Tau in neuronal cell bodies/ somata treated with non-toxic doses of α-IgLON5#1 or pCtrl (0.1, 1, 10 μg/ml) for 2 days. N = 3 experiments with 3-5 replicates. Data points show individual neurons (22-95 neurons per group). (**C**) Total Tau in cell bodies upon treatment with 1 μg/ml of α-IgLON5 AABs from four different patients or pCtrl for 2 days. N = 3 experiments with 3-5 replicates. Data points show individual neurons (43-136 neurons per group). (**D**) Total Tau in cell bodies upon treatment with 1 μg/ml of α-IgLON5#1or pCtrl for 5 days continuously (bars without pattern fill) or for 2 days followed by 3 days without AABs (bars with pattern fill). N = 3 experiments with 3-5 replicates. Data points show individual neurons (48-71 neurons per group). **(B, C, D)** One-way ANOVA with Tukey post-test.

Importantly, Tau accumulation in neuronal cell bodies also occurred upon treatment with AABs from other anti-IgLON5 patients, α-IgLON5#2 and α-IgLON5#4, that also showed strong binding to the neuronal surface (Fig. 1C), however, not for the weaker binding α-IgLON5#3 (Fig. 2C; fig. S2F). Together, these data suggested that binding of IgLON5 AABs can directly trigger Tau missorting. Interestingly, when removing α-IgLON5 AABs from the medium after 2 days and quantifying cell body Tau immunofluorescence 3 days later, the cell body Tau levels remained similar to conditions of continuous treatment with AABs (Fig. 2D), suggesting that Tau, once resorted, resides in the somatodendritic compartment for multiple days. Notably, by Western blot, overall total and phosphorylated Tau (AT8, PHF-1, AT180, pS199 epitopes) were similar in whole cell lysates of α-IgLON5 AAB-, pCtrl-, and non-treated neurons (fig. S2G, H). This was likely due to generally high Tau phosphorylation in cultured neurons at baseline (*27*).

### α-IgLON5 AABs induce hippocampal Tau phosphorylation

Next, we assessed whether α-IgLON5 AABs would be capable and sufficient of inducing Tau changes *in vivo*. For this, we delivered α-IgLON5 AABs into the brains of adult wildtype mice, which usually do not develop Tau aggregation pathology but can show abnormal Tau phosphorylation in sporadic tauopathy-associated conditions (e.g., seizures (*28*, *29*) or traumatic brain injury (*30*)). Over the course of 14 days, a total amount of 75 μg (∼3 μg/g body weight) of α-IgLON5#1 was infused via Alzed pumps into the right lateral ventricle (N = 10 animals). Littermates infused with the same amount and concentration of pCtrl (N = 9), or with PBS (N = 9), functioned as controls. After the infusion, the brains were analyzed for phospho-Tau and neuroinflammation by immunohistology, and RNA and protein were extracted from the contralateral hemisphere for transcriptome and protein analyses (Fig. 3A).

**Fig. 3.**
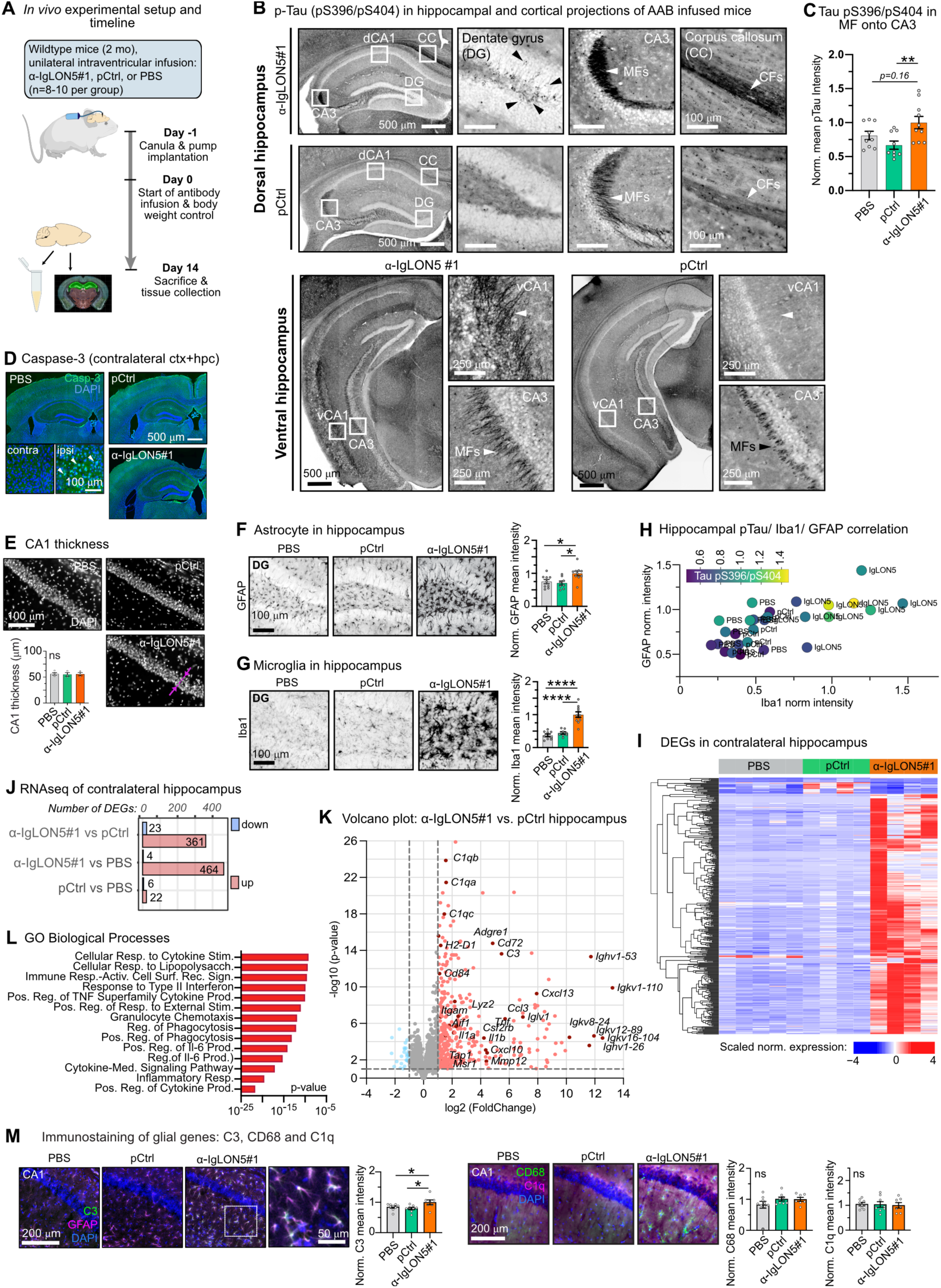
Intraventricular α-IgLON5 AAB infusion triggers hippocampal Tau phosphorylation and glia responses. (**A**) Experimental design for *in vivo* application of α-IgLON5 AABs. Wildtype mice were infused in the right ventricle with α-IgLON5#1, pCtrl, or PBS. 75 μg AABs were infused over 14 consecutive days. At Day 14, serum samples were collected before animals were sacrificed. Brain tissue was dissected and then fresh-frozen or PFA-fixed for analysis of protein and RNA content or by immunofluorescence. (**B**) Representative images of p-Tau pS396/pS404 immunoreactivity in α-IgLON5#1 and pCtrl mouse brains. phospho-Tau accumulation is observed in dorsal hippocampus (top panel) in dentate gyrus (DG) granule cells (black arrowheads), mossy fiber (MF) projections onto CA3 (white arrowhead), and commissural fiber (CF) tracks of the corpus callosum (CC; white arrowhead). In the ventral hippocampus, phospho-Tau accumulation is found in projections onto ventral CA1 (vCA1; white arrowhead) and MF projections onto CA3 (white arrowhead). (**C**) Quantification of Tau pS396/pS404 intensity in projections onto CA3. N = 8-10 animals per group, 3-5 brain sections per animal. (**D**) Immunolabeling of Caspase-3 in α-IgLON5#1 and control brains. Caspase-3 positive cells (white arrow heads) are only found in the ipsilateral cortex near the injection site. Scale bars as indicated. (**E**) Images of hippocampal CA1 pyramidal layer in α-IgLON5#1 and control brains, with quantification of CA1 layer thickness (measured as indicated by pink line with arrow heads). N = 3-4 animals, 3-4 brain sections per animal. (**F**) Representative images of astrocytes (GFAP) in DG of α-IgLON5#1 and control animals, with quantification of GFAP intensity in entire hippocampal regions of α-IgLON5#1 and control brains. N = 8-10 animals, 3-5 brain sections per animal. (**G**) Representative images of microglia (Iba1) in DG of α-IgLON5#1 and control animals, with quantification of GFAP intensity in entire hippocampal regions of α-IgLON5#1 and control brains. N = 8-10 animals, 3-5 brain sections per animal. (**H**) Triple correlation of hippocampal GFAP vs. Iba and Tau pS396/pS404 (color gradient) fluorescence intensity levels. Data points represent individual animals. (**I**) Hierarchical clustered heatmap of differentially expressed genes (DEGs) in contralateral hippocampi of PBS, pCtrl, and α-IgLON5#1 infused mice. (**J**) Number of significantly upregulated (red) and downregulated (blue) DEGs in contralateral hippocampi of PBS, pCtrl, and α-IgLON5#1 animals. (**K**) Volcano plot of gene deregulation in contralateral hippocampi of α-IgLON5#1 compared to pCtrl animals. (**L**) Gene Ontology (GO Biological processes) enrichment analysis of upregulated genes in hippocami of α-IgLON5#1 compared to pCtrl animals. (**M**) Immunolabeling of neuroinflammatory markers (C3, CD68, C1q) in hippocampi of α-IgLON5#1 and control mice. For the quantification of C3, CD68 and C1q fluorescence intensity, data are shown as mean±SEM, N = 7-9 animals, 3-5 brain sections per animal. **(C, E, F, G, M)** Data in shown as mean±SEM. One-way ANOVA with Tukey post-test. **(B, D, E, F, M)** Scale bars as indicated.

Immunodetection of α-IgLON5#1 showed strong labeling in brain areas around the infused ventricle (cortex, hippocampus, striatum), where the AAB concentration is expected to be the highest, and weaker labeling in more distant ipsilateral regions (e.g., parts of thalamus, hypothalamus, amygdala) (fig. S3A). In the contralateral hemisphere, mostly the hippocampus showed α-IgLON5#1 immunoreactivity. Animals treated with pCtrl, although having received the same antibody dose, showed generally less labeling, also in the contralateral hippocampus. This is in line with the weak, unspecific binding of pCtrl observed in neuronal cultures. In the following, we analyzed only tissue from contralateral hemispheres to avoid artifacts induced by injury and scar formation associated with the canula implantation. Immunolabeling of brain sections revealed higher levels of phosphorylated Tau-pS396/pS404 in the contralateral hippocampus of α-IgLON5#1 compared to control (pCtrl and PBS) mice, particularly in mossy fiber projections onto CA3 (IgLON5#1 vs. pCtrl: *p* = 0.01). Tau-pS396/pS404 could also be found in some cell bodies in dentate gyrus, in bi-directional projections on ventral (but not dorsal) CA1, and in commissural fibers of the corpus callosum (Fig. 3B, C). In other brain regions (including spinal cord, fimbria, cerebellum, and hypothalamus), we did not observe Tau-pS396/pS404 immunoreactivity. For Tau-pS202/pT205 (AT8 epitope), we observed no immunoreactivity in hippocampal or other brain areas (fig. S3B). Notably, detecting Tau changes in wildtype mice – normally rather resilient to pathological changes in endogenous mouse Tau – demonstrates the strong potential of α-IgLON5 AABs in triggering Tau changes.

Next, we examined whether the increased Tau phosphorylation induced by α-IgLON5 AAB infusion was associated with neurotoxicity. However, we found no signs of apoptosis in the hippocampus, assessed based on the absence of Caspase-3 positive cells (Fig. 3D) as well as immunolabeling of milk fat globule epidermal growth factor 8 (MFG-E8 (*31*)) (fig. S3C). Caspase-3 positive cells were only present in the ipsilateral cortex near the injection site due to tissue damage during pump implantation (Fig. 3D). We also did not observe neurodegeneration, assessed based on CA1 neuronal layer thickness (Fig. 3E) and neurofilament light chain (Nfl) serum levels (*32*) (fig. S3D). Together these data indicated that 2 weeks of cerebroventricular infusion of α-IgLON5 AABs induced local, hippocampal Tau phosphorylation in the absence of detectable neurotoxicity. However, α-IgLON5#1 animals had slightly lower body weight (non-significant) compared to control groups throughout the infusion period (fig. S3E).

### Neuroinflammation caused by α-IgLON5 AABs

Tau phosphorylation and missorting, prior to aggregation and neurodegeneration, were previously suggested to correlate with glia cell activation in tauopathy animal models and human brains (*33*, *34*). Similarly, we found that astrocytic GFAP (Fig. 3F; fig. S3F) and microglial Iba1 intensities (Fig. 3G; fig. S3G) were significantly increased in the contralateral hippocampus of α-IgLON5#1-infused mice compared to pCtrl and PBS controls. On the animal level, hippocampal Iba1 and GFAP levels appeared to increase with mossy fiber Tau-pS396/pS404 intensity, based on linear regression (moderate correlation by spearman) (Fig. 3H; fig. S3H).

To gain further insights about α-IgLON5#1-induced effects in the brain, we analyzed the gene expression (by RNA-Seq) in contralateral hippocampi and cortices. In hippocampi of α-IgLON5#1, we detected 23 significantly downregulated and 361 significantly upregulated transcripts compared to pCtrl animals (Fig. 3I, J; fig. S3I). Most of these transcripts were associated with inflammatory pathways (GO Biological Processes) particularly IL-6 and TNFα cytokine responses, or belonged to immunoglobulin genes (Fig. 3K, L; Data S1, S2). The cortical transcriptome of the mice showed similar changes between α-IgLON5#1 and pCtrl animals, while only few differences in gene expression were found between the two control groups, pCtrl and PBS (fig. S3J, K). Of the genes upregulated in the hippocampus of α-IgLON5#1 animals (Fig. 3I-K;), we confirmed the upregulation of the neuroinflammation-associated astrocyte gene complement-3, C3, by immunolabeling (Fig. 3M).

Together the *in vivo* data show that cerebroventricular infusion of α-IgLON5 AABs induces localized hippocampal epitope-specific (pS396/pS404) Tau phosphorylation and neuroinflammation in wildtype mice. In contrast to the animal model used here, in which mice received a direct AAB infusion selectively into the right ventricle for 2 weeks, anti-IgLON5 disease patients’ brains are diffusely exposed to the AAB over prolonged times. Such differences in AAB exposure likely explain the different distribution of Tau changes and neuroinflammation in humans, where Tau phosphorylation is mostly found in brain stem, hypothalamus and cerebellum (*3*, *7*, *8*, *22*) (Supplementary Text).

### α-IgLON5 AABs trigger acute neuronal hyperactivity

Somatodendritic Tau missorting has previously been suggested to be driven by neuronal hyperactivity, for example, in the context of epileptic activity (*35*), Aβ oligomers (*36*), and chemical stimulation, e.g. glutamate (*36*). Since α-IgLON5 AABs induced Tau missorting, we hypothesized that α-IgLON5 AABs may exert neuronal hyperactivity.

To test this hypothesis, we assessed spontaneous neuronal activity using Calcium (Ca^2+^) imaging in gCamp6f-expressing neurons (Fig. 4A) treated with α-IgLON5 AABs or pCtrl (1 μg/ml). One hour after treatment, a significant increase in Ca^2+^ spike frequency could be detected for patient-derived AABs α-IgLON5#1, α-IgLON5#2, and α-IgLON5#4 compared to pCtrl or non-treated (NT) neurons (Fig. 4B, C). α-IgLON5#3, which showed inefficient binding and did not induce Tau missorting (Fig. 1C, 2C), also did not increase neuronal activity. Activity-dependent expression of the immediate early gene c-FOS confirmed the observed increase in activity, showing significantly increased fluorescence intensity of nuclear c-FOS in α-IgLON5#1 and α-IgLON5#2 treated neurons (Fig. 4D). Even at a low AAB dose (0.1 μg/ml ∼ 0.67 nM), at which α-IgLON5#1 binding to neurons was similar to pCtrl (Fig. 1C), α-IgLON5#1 still induced a significant increase in neuronal activity compared to pCtrl (Fig. S4A-C). The mild activity induced by pCtrl in this condition can be accounted to the antibody application procedure, since a non-binding human monoclonal antibody (mCtrl (*37*)) showed a similar effect. In summary, these data indicated that α-IgLON5 AAB binding correlated with neuronal activity and Tau missorting (fig. S4D, E). Importantly, application of EGTA-AM to suppress intracellular Ca^2+^ spikes (*38*) prevented somatodendritic Tau missorting (Fig. 4E,F; fig. S4F) thereby establishing a causal link between α-IgLON5 AAB-induced hyperactivity and Tau missorting. Furthermore, also the treatment of neurons with the sodium channels blocker tetrodotoxin (TTX) prior to α-IgLON5#1 treatment abolished α-IgLON5#1 induced hyperactivity (Fig. 4G). This could indicate that Ca^2+^ and sodium channels may be involved in α-IgLON5 AAB-induced activity.

**Fig. 4.**
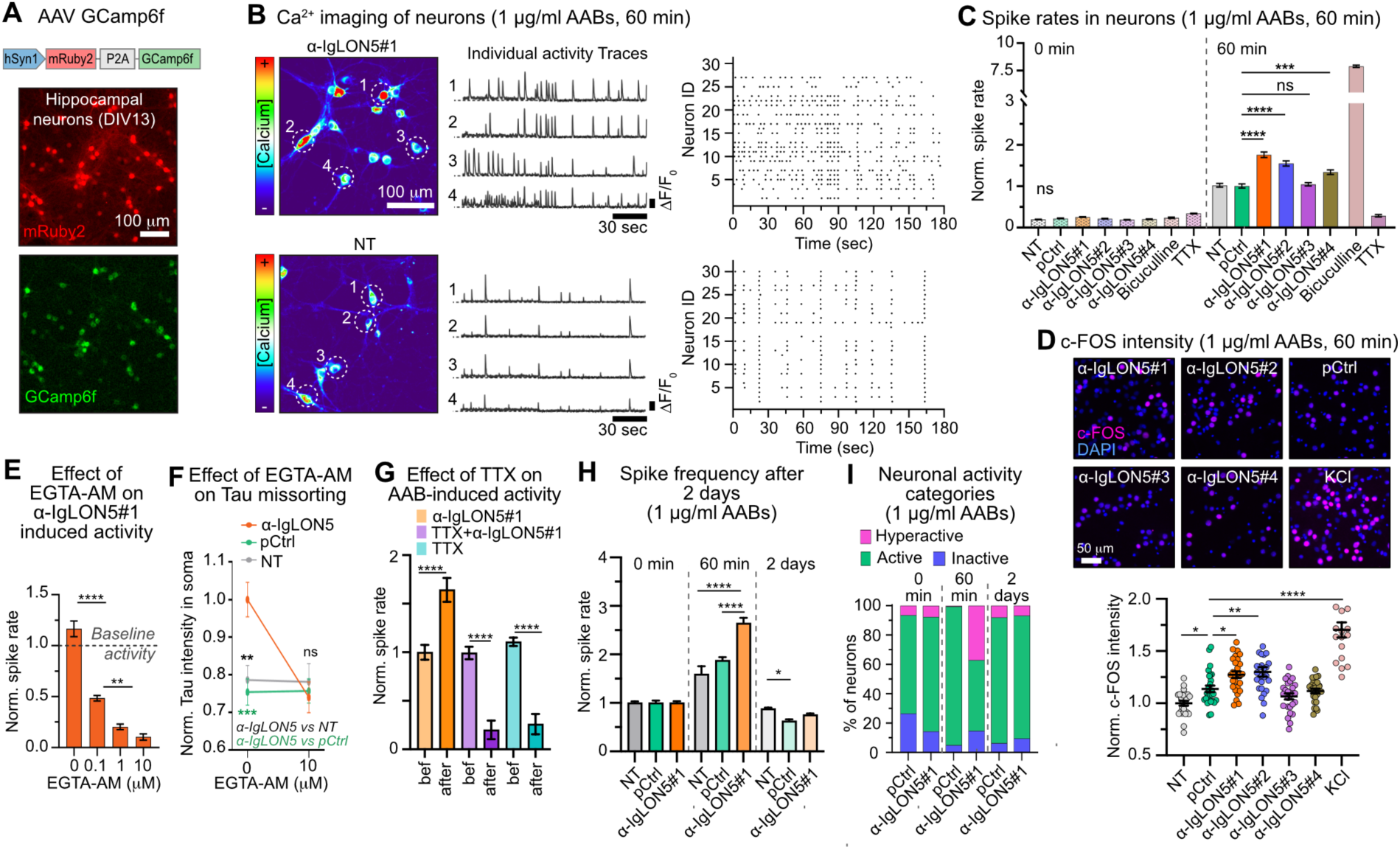
α-IgLON5 AABs induce neuronal hyperactivity. (**A**) Expression (by AAV) of mRuby2 (transduction marker) and Ca^+2^ indicator GCamp6f under hSyn1 promoter in cultured neurons. (**B**) Ca^2+^ levels (time-integrated; pseudo-colored) from time-lapse recordings of GCamp6f+ neurons treated with 1 μg/ml α-IgLON5#1, or no-treatment control (NT). White ROIs indicate positions of neurons for shown GCamp6f intensity traces (ΔF/F_0_ vs time). Raster plots show occurrence of individual Ca^+2^ spikes in cell bodies of neurons in one field of view (N = 25-30 cells, y-axis). (**C**) Normalized Ca²⁺ spike rates in neurons treated with 1 μg/ml of different patient-derived α-IgLON5 AABs (α-IgLON5#1, #2, #3, and #4) or pCtrl for 60 min. Ca^+2^ spikes were recorded before (at 0 min) and after (at 60 min) AAB treatment in the same cultures. Pharmacological controls: Bicuculline (30 μM for 5 min) induced hyperactivity, TTX (0.5 μM for 10 min) silenced neurons. N = 3 experiments, 875-1344 neurons per group. Data normalized to untreated neurons at 60 min treatment. **D**) Images of c-FOS immunostaining in neurons treated with 1 μg/ml α-IgLON5 AABs or pCtrl for 60 min. KCl treatment (8 mM, 60 min) as positive control. N = 3 experiments. (**E**) Ca^+2^ spikes in neurons pretreated with different EGTA-AM concentrations for 20 min, then exposed to α-IgLON5#1 (1 μg/ml) for 60 min. Data normalized to untreated neurons. N=3 experiments, 203-309 neurons per condition. (G) Tau missorting (cell bodies intensity) in neurons pretreated or not with 20 μM EGTA-AM for 20 min, then exposed to 1 μg/ml α-IgLON5#1 or pCtrl for 2 days. N = 3 experiments, 47-60 neurons per conditions. (**G**) Normalized spike rates before and after treatment with α-IgLON5#1 (1 μg/ml for 60 min), TTX (0.5 μM for 10 min), or their combination (0.5 μM TTX for 10 min, then 1 μg/ml α-IgLON5#1 for 60 min). N = 3 experiments. (**H**) Ca^+2^ spikes in neurons before (0 min) and after 1 μg/ml α-IgLON5#1 or pCtrl treatment for 60 min or 2 days. N = 3-4 experiments, 263-309 neurons per conditions. (**I**) Percentage of inactive (blue), active (green), and hyperactive (pink) neurons in cultures treated with 1 μg/ml α-IgLON5#1 or pCtrl for 0 min, 60 min, or 2 days. Hyperactivity criteria: spikes/min > mean + 2*SD of pCtrl at each recording time point. Active: 0 < spikes/min < mean+2*SD of pCtrl at each recording time point. Silent: no spikes. Panels (**C-H)**: Data shown as mean ± SEM, One-way ANOVA with Tukey post-test. **(A, B, D)**: Scale bars as indicated.

Next, we investigated how α-IgLON5 AAB or pCtrl (1 μg/ml) exposure for 2-3 days would impact neuronal activity, a time point when Tau was missorted into neuronal cell bodies following α-IgLON5 AAB treatment. After the initial induction of neuronal hyperactivity within 60 min after α-IgLON5#1 application, both the average spike rate and the fraction of hyperactive neurons dropped back to levels detected before treatment (Fig. 4H, I; fig. S4G). This was confirmed by electrophysiological recordings in autaptic cultures of hippocampal neurons (fig. S4H, I) treated with AABs for 3 days, which showed no difference between α-IgLON5#1 and pCtrl-treated cells for any measured parameters (fig. S4I). Furthermore, activity-dependent c-FOS expression in the DG was also similar in mice that had received α-IgLON5#1 or pCtrl antibodies for two weeks (fig. S4J). α-IgLON5 AAB-induced neuronal hyperactivity therefore seemed to be transient, whereas the related Tau missorting persisted over days.

Prolonged AAB exposure may lead to a reduction of synapse numbers, which may dampen the initial α-IgLON5 AABs-induced hyperexcitation. Previous data from neurons treated with high doses of antibodies from anti-IgLON5 disease patient serum (1:50 dilution of total IgG isolates) showed a loss of synaptic protein content indicative of synaptic decline (*21*). In our model of low dose (1 μg/ml) α-IgLON5#1 treatment, however, no significant loss in pre- and postsynaptic marker densities (fig. S4K, L) was observed compared to control groups up to 7 days of AAB treatment.

### α-IgLON5 AABs cluster cell adhesion and ion channel proteins

The decline of hyperactivity after multiple day of α-IgLON5 AAB treatment could be due to internalization of IgLON5 protein/antibody complexes, leading to a decrease in IgLON5 cell surface epitopes (*2*, *19*, *39*). Previous studies reported a loss of surface IgLON5 clusters in neurons after 3 days of treatment with the bulk antibody fraction from α-IgLON5 patient serum (*19*). In neurons continuously treated with α-IgLON5#1 over multiple days, we found an initial increase in IgLON5 surface clusters compared to controls after 60 min, followed by a decline of IgLON5 clusters back to densities comparable to untreated control neurons after 7 days (Fig. 5A). We qualitatively confirmed internalization of pHrodo-labeled α-IgLON5#1 after 2 days, and to a lesser amount also for pCtrl ABs, which may be due to unspecific antibody/protein internalization by neurons (fig. S4M). IgLON5 cluster internalization could therefore in part account for the observed reduction in hyperactivity after 2-3 days. After 60 min of α-IgLON5#1 treatment, however, only ∼7.5% of IgLON5 immunofluorescence signal was intracellular (fig. S4N).

**Fig. 5.**
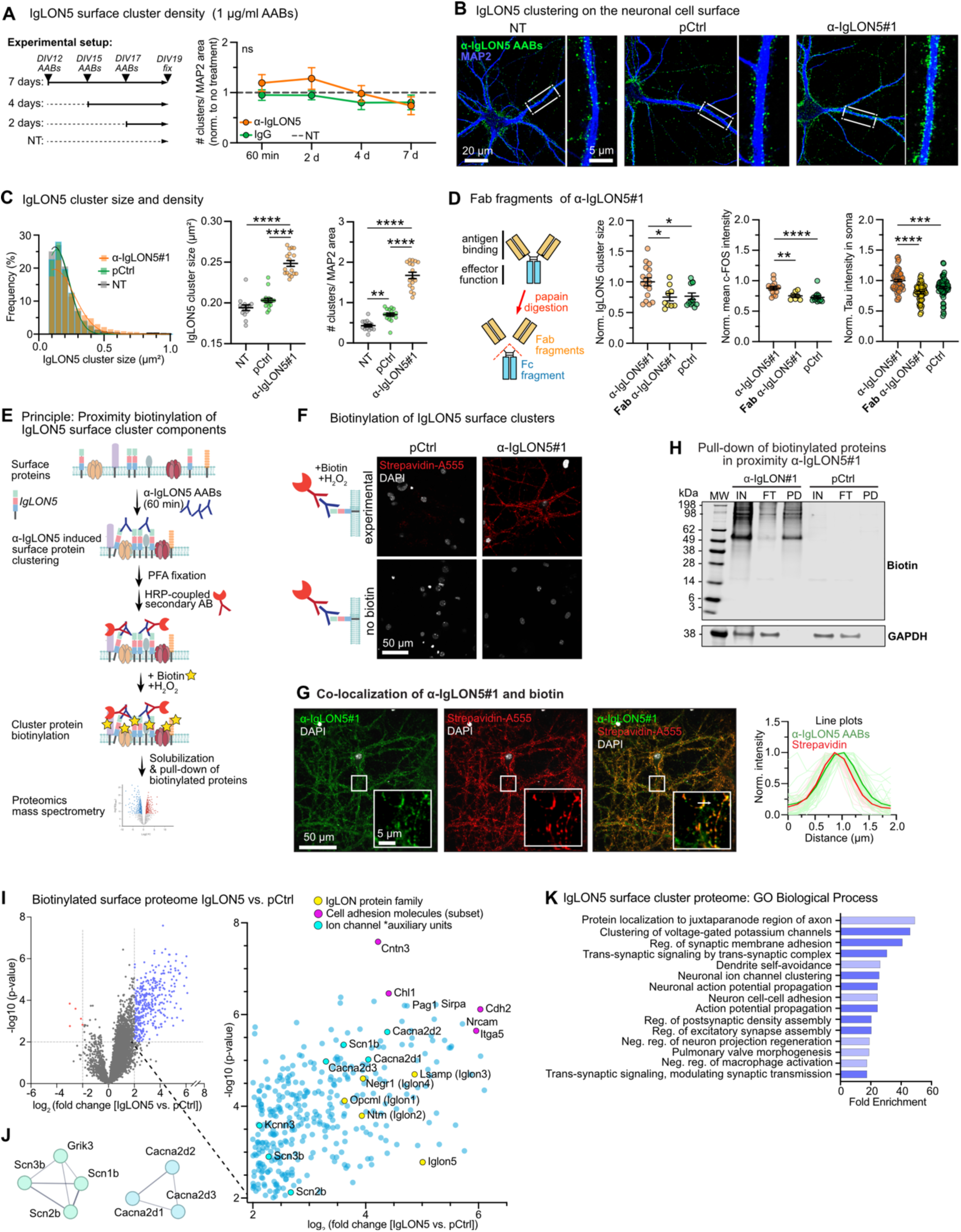
α-IgLON5 AAB-induced clustering of cell surface proteins. (**A**) Quantification of IgLON5 surface cluster density (# clusters/MAP+ area) in primary neurons treated with 1 μg/ml α-IgLON5#1 or pCtrl at indicated time points (60 min, 2, 4, and 7 days). N = 3 experiments with 3-4 replicates per condition. Two-way ANOVA with Sidak post-test. (**B**) Confocal images of IgLON5 surface clusters on non-permeabilized neurons upon treatment with 1 μg/ml α-IgLON5#1 or pCtrl for 60 min. Scale bars as indicated. (**C**) Refined analysis of IgLON5 surface cluster density and size. Histogram shows frequency distribution of IgLON5 surface cluster size for all measured clusters. Additionally, mean cluster size and density (#IgLON5 clusters/ MAP+ area) are shown. N = 3 experiments with 3 replicates per condition. One-way ANOVA with Tukey post-test. (**D**) α-IgLON5#1 and pCtrl Fab fragment (antigen-binding region) generation by papain digestion. IgLON5 surface cluster size/area, neuronal activity (measured as nuclear c-FOS intensity), and Tau accumulation in cell bodies in neurons treated with equimolecular concentrations of α-IgLON5#1, Fab α-IgLON5#1 and pCtrl. For analysis of cluster size and neuronal activity, neurons were treated for 60 min. For Tau missorting, neurons were treated for 2 days. N = 3 experiments with 4-5 replicates per condition. One-way ANOVA with Tukey post-test. (**E**) Antibody-mediated proximity biotinylation assay. Live hippocampal neurons were treated with α-IgLON5#1 for 60 min to allow surface clustering, then fixed and incubated with α-human IgG-HRP. Membrane-impermeant biotin-phenol and H₂O₂ were applied to selectively label surface proteins in proximity to AAB-bound IgLON5 clusters. Biotinylated proteins were pulled-down and analyzed by mass spectrometry. Surface protein biotinylation could be observed based on streptavidin-Alexa555 labeling in neurons treated with α-IgLON5#1, α-human IgG-HRP, and biotin. (**F**) Detection of IgLON5 surface cluster biotinylation with streptavidin-Alexa555 upon incubation of neurons with 1 μg/ml α-IgLON5#1, followed by fixation, HRP-secondary incubation and biotinylation reaction. Scale bar as indicated. (H) Representative images of overlapping α-IgLON5 and Streptavidin-Alexa555 signal in neurons treated with 1 μg/ml α-IgLON5#1 for 60 min. Intensity profile plot confirms co-localization of α-IgLON5 AABs and biotin signal. Scale bars as indicated. (**H**) Example western blot showing biotinylated proteins in input (IN = cell lysate), flow-through (FT = wash), and pulldown (PD). Note, GAPDH is only detected in IN and FT samples, confirming cell surface specificity of the labeling. (**I**) Volcano plot of biotinylated surface proteins in neurons treated with α-IgLON5#1 versus pCtrl. Proteins significantly enriched in IgLON5 clusters (blue points) are provided as magnified view: IgLON family members (yellow; IgLON51/2/3/4/5), cell adhesion proteins (pink), and ion channel subunits (cyan). (**J**) Interaction network (STRING) of selected ion channel auxiliary subunits found in IgLON5 clusters (from Fig. S5E). (**J**) Gene Ontology (GO) analysis of biological processes associated with proteins significantly enriched in α-IgLON5 AAB-induced surface clusters.

AAB-induced clustering of surface receptors, i.e., NMDARs and LGI1, can change receptor surface availability and dynamics, leading to altered neuronal activity (*40–46*). We therefore hypothesized that AAB-induced IgLON5 clustering may be involved in neuronal hyperactivity as well. Untreated cultured neurons showed a clustered appearance of IgLON5 on the cell surface (Fig 5B (*19*)). Acute (60 min) treatment with α-IgLON5 AABs upregulated IgLON5 surface cluster size and their dendritic density (Fig. 5A-C). This occurred at maintained total cell surface labeling of IgLON5, indicating no increase in IgLON5 protein content (fig. S5A)

To test whether α-IgLON5 AABs directly induced the clustering, we separated the antigen-recognizing Fab fragments from the connecting Fc part of α-IgLON5#1 by papain digestion (fig. S5B), which removes the ability of the AAB to bind and “crosslink” multiple IgLON5 molecules at a time. This approach was previously used to disable clustering of NMDARs by α-NMDAR AABs (*47*). For α-IgLON5#1, treatment with equimolar concentrations of α-IgLON5#1 Fab fragments (2 mol Fab fragments for 1 mol IgG) did not induce IgLON5 surface clustering as observed for intact α-IgLON5#1 (Fig. 5D; fig. S5C). α-IgLON5#1 Fab fragments also induced less neuronal activity (c-FOS intensity) and Tau missorting (Fig. 5D). Hence, α-IgLON5#1-induced IgLON5 clustering seemed to be involved in inducing neuronal hyperactivity and Tau changes.

To determine whether IgLON5 clusters contained surface molecules capable of inducing neuronal hyperactivity, we determined the IgLON5 cluster proteome using an immunolabeling-based proximity biotinylation approach (Fig. 5E, F; fig. S5D). Neurons treated with α-IgLON5#1 or pCtrl for 60 min were fixed (4%PFA in PBS) and immunolabeled with α-human IgG secondary coupled to horseradish peroxidase (HRP). Addition of cell-impermeable biotin and hydrogen peroxide (H_2_O_2_) catalyzed the biotinylation of cell surface proteins in proximity to IgLON5#1, hence, in IgLON5 cell surface clusters. By microscopy, the spatial overlap and restriction of biotinylation to IgLON5 clusters could be confirmed (Fig. 5G). We therefore considered biotinylated proteins in α-IgLON5#1-treated neurons to be part of IgLON5 clusters. Western blots confirmed the biotinylation of proteins in lysates of α-IgLON5#1-treated neurons, and to a lesser degree in pCtrl-treated neurons (Fig. 5H), and the successful pull-down of these proteins using streptavidin beads. Proteomics mass spectrometry revealed the enrichment of specific cell surface proteins in IgLON5 clusters: other members of the IgLON family (IgLON1/2/3/4) – as expected from a previous study (*15*) –, as well as other cell adhesion molecules (i.e., Ncam, Nrcam, Cadherins (Cdh4, Cdh6, Cdh11, Cdh2), Contactins (Cntn1-6) and Integrins (Itga5, Itga7, Itgb1); Fig. 5I, Data S3). IgLON5 clusters also contained ion channel auxiliary units involved in the regulation of neuronal activity, in particularly regulatory subunits of voltage-gated sodium channels (Scn1b/2b/3b) and voltage-gated Ca^2+^ channels (Cacna2d1/d2/d3), voltage gated potassium channel KCNN3, and the kainate receptor subunit GRIK3 (Fig. 5J; fig. S5E). Notably, we did not detect Na_v_ and Ca_v_ in the IgLON5 clusters proteome. This may be attributed to their embedding in the plasma membrane, which renders the channels difficult to extract from fixed neurons and may make them inaccessible for the biotinylation reaction using non-cell permeable biotin. Gene ontology analysis (GO Biological Processes) of all differentially enriched proteins revealed proteins involved in cell adhesion, synapse assembly and function, synaptic and axonal signal transmission and propagation, and ion channel clustering (Fig. 5K; fig. S5F; Data S4). Importantly, the proteome of the input cell lysate (prior to streptavidin bead pull-down) of α-IgLON5#1 and pCtrl treated neurons did not have significant differences (fig. S5G, Data S3).

In summary, our data suggest that the binding of α-IgLON5 AABs induces physical clustering and stimulation of Na_v,_ Ca_v_, and K_v_ channels and auxiliary subunits and the glutamate-sensing kainate receptor subunit GRIK3. The deregulation of this complex mix of activity regulating channels and receptors by α-IgLON5 AABs explains the acute and excessive hyperexcitability of neurons upon α-IgLON5 AABs binding, which ultimately triggers prolonged somatodendritic Tau missorting and neurotoxicity. In addition, α-IgLON5 AABs induce clustering of cell adhesion molecules, including cadherins and integrins, a fundamental step at the top of cell adhesion signaling cascades. Whether cell adhesion signaling contributes to α-IgLON5 AAB induced Tau changes has to be further investigated.

## Discussion

AABs against neuronal receptors and cell surface proteins, e.g., NMDAR, GABA_A_R, and LGI1, can alter neuronal physiology (*1*, *2*, *48–50*). Our data reveal that anti-IgLON5 disease may involve a similar disease mechanism: cell surface binding of α-IgLON5 AABs promotes the physical clustering of surface IgLON5 with cell adhesion molecules and ion channels, leading to acute neuronal hyperactivity that induces persistent Tau missorting, promotes Tau aggregation, and ultimately can lead to neurotoxicity. Previous studies reported that long-term treatment (for weeks) of neurons *in vitro* or of mice with anti-IgLON5 disease patient serum antibody pools (not enriched for α-IgLON5 AABs) disrupted synaptic protein levels, neuronal activity, and caused cognitive deficits and anxiety-like behavior in mice (*21*, *51*, *52*). Our data now sheds light on the events upstream of these effects.

Antibody-induced changes in receptor surface distribution have previously been reported for AABs. For example, AABs against LGI1 reduce its surface availability and impair interactions with potassium channels, leading to destabilization of synaptic contacts, dysregulation of excitatory transmission, and initiation of pathological hyperactivity including seizures (*43*, *44*, *46*, *50*, *53*). α-NMDAR AABs trigger nanoscale clustering of NMDARs, disrupting their association with scaffolding proteins, followed by receptor internalization and degradation, ultimately leading to a reduction in surface NMDARs and impaired synaptic transmission and neuronal silencing (*41*, *42*, *47*, *49*). For α-IgLON5 AABs, we similarly found that binding induces clustering of IgLON5 together with cell surface proteins. After 2 days, however, the enhanced IgLON5 receptor/protein clustering disappeared, likely due to internalization. If α-IgLON5 AAB induced hyperexcitation is directly related to IgLON5 clustering – as suggested by our data – the decline in surface clusters can explain why AAB-induced hyperexcitation vanished after 2 days in neuronal cultures.

α-IgLON5 AABs induce clustering of ion channel auxiliary units (Scn1b/2b/3b, Cacna2d1/d2/d3) that regulate the permeability of voltage-gated Ca^2+^ (Ca_v_) and sodium (Na_v_) channels, which are essential for synaptic transmission and action potential initiation and propagation (*54*), and with the Ca^2+^-activated potassium channel KCNN3 and the ionotropic glutamate receptor GRIK3/GLUR7, which are directly involved in neuronal excitability (*55*, *56*). Aberrant physical interactions and clustering of these proteins – induced by α-IgLON5 AABs - can have a profound impact on neuronal activity. In addition, we identified several cell adhesion molecules (IgLON family members, integrins, cadherins, protocadherins, contactins) in IgLON5 surface clusters. These transmembrane proteins dynamically interact with the actin cytoskeleton to facilitate synaptic organization, neuronal migration, and development (*57*, *58*). Their interaction with other surface proteins modulates function and localization of voltage-gated ion channels, thereby altering Ca^2+^ influx, neuronal excitability, and other signaling pathways that remodel the cytoskeleton, strengthen adhesion, support synaptic function and plasticity. The concerted effect of α-IgLON5 AAB binding on adhesion proteins and ion channels explains how α-IgLON5 AAB binding induces neuronal hyperexcitability. Whether individual molecular players are most relevant for inducing neuronal hyperactivity upon α-IgLON5 AAB binding, and which downstream signalling cascades are involved in Tau activation, needs to be clarified.

Different patient-derived α-IgLON5 AABs triggered non-toxic, acute (minutes to hours) neuronal hyperactivity, which was linked to persistent (multi-day) somatodendritic Tau missorting and promotion of Tau aggregation. In the brain of anti-IgLON5 disease patients, this could allow for a gradual, irreversible and neurotoxic accumulation of pathological Tau. The extended presence of Tau in the somatodendritic compartment can exhaust neuronal homeostasis (*59*), induce synapse loss (*60*), and impair other key neuronal functions (*61–63*). Consequently, this would explain why immunotherapy (to deplete AABs) seems most beneficial when initiated early in anti-IgLON5 disease (*5*), i.e., prior to overt Tau accumulation. Importantly, Tau pathology has previously been connected to neuronal hyperactivity in epilepsy (*64*) and in AD (*65–67*). Susceptibility of neurons to hyperexcitation could therefore encode a selective vulnerability of neuronal circuits to Tau pathology. The here revealed hyperactivity-induced Tau pathology in anti-IgLON5 disease emphasizes the importance of considering the modulation of neuronal activity for therapeutic approaches to prevent or lower Tau pathology and toxicity across tauopathies. For the treatment of anti-IgLON5 disease patients, determining the neuromodulatory activity of their plasma AAB pool could help in the choice of treatment, i.e., whether to target neuronal hyperexcitation, neuroinflammation, Tau phosphorylation and aggregation, or all. For example, our data indicate that the effects of α-IgLON5 AABs on neuronal activity and Tau depend on AAB binding strength, AAB concentration, and duration of exposure. These factors differ between patients and may contribute to variations in brain functional impairment and Tau pathology progression between anti-IgLON5 patients (*8*). Further, whether neuronal hyperactivity plays a role for Tau pathology in other immune-related secondary tauopathies, e.g., in subacute sclerosing pan-encephalitis (SSPE (*68*); measles late reaction) and head-nodding syndrome (*69*) (involving DJ-1 antibodies), should be tested as well.

In mice with cerebroventricular infusion of α-IgLON5 AABs, Tau phosphorylation occurred in hippocampal projections (i.e., mossy fibres between dentate gyrus and hippocampal CA3, projections onto ventral CA1, and commissural fibres) and was accompanied by microglia activation and a neuroinflammatory gene expression signature (TNFα and IL-6 pathways). These changes are reminiscent of early pathological alterations found in other tauopathies, like AD and FTD, where changes in Tau (i.e., phosphorylation and missorting) spatially associate with progressive decay in synaptic homeostasis (*60*, *70–72*) and neuroinflammation and, ultimately, lead to neurodegeneration (*8*, *9*) (Supplemental Text). In addition, α-IgLON5 AABs appear to trigger a pronounced upregulation of Ig gene expression typically not reported in tauopathy brains or models. This may indicate B-cell infiltration, e.g., as a result of α-IgLON5 AAB induced IL-6 upregulation (*73*), or inflammation-related microglial Ig expression (*74*). Mild B cell infiltration has been reported in IgLON5 autopsy brains (*22*). Whether Ig gene upregulation occurs in IgLON5 disease patient brains is not known. The IgG subclass composition of α-IgLON5 AAB in the brain may play an important role for the extend and kind of neuroinflammation induced (Supplemental Text). As in other tauopathies, however, it remains uncertain whether α-IgLON5 AAB-induced neuroinflammation is mediated by Tau changes and to which extend neuroinflammation contributes to Tau pathology.

## Materials and Methods

### Ethics

All analyses of patient material were approved by the Charité - Universitätsmedizin Berlin Ethics Board (#EA1/258/18). Primary mouse neuron preparations and experiments involving adult animals (C57/B6, male) were carried out according to the guidelines stated in the directive 2010/63/EU of the European Parliament on the protection of animals used for scientific purposes and were approved by local authorities in Berlin and the animal welfare committee of the Charité Universitätsmedizin Berlin, Germany.

### IgLON5 patients

Anti-IgLON5 disease patients, whose α-IgLON5 ABB and pCtrl AB pools were isolated in this study, were patients in the neurology clinics of the Charité University Medicine Berlin (patients: IgLON5#2,3,4) and of the Otto-von-Guericke-University Magdeburg (patient: IgLON5#1) (Table 1). Phosphorylated tau 217 (pTau217) in serum of patients IgLON5#2,3,4 and in plasmapherisate of patient IgLON5#1 were measured using the Simoa® pTau217 Advantage PLUS assay (Quanterix, Billerica, MA, USA; kit lot 504407) at the DZNE Bonn. The intra-plate CV was <5.1% based on two control levels run in duplicates at the beginning and end of the plate.

**Table 1.** Anti-IgLON5 disease patient information. MP = Methylprednisolone, PPH = plasmapheresis, RTX = Rituximab, IA = Immunoadsorption, IVIG = intravenous immunoglobulins, * = pTau217 in plasmapherisate of this patient.

| Patient# | Sex (M/F) | Age (Y) | Immuno-therapy | Effect? | AAB titer | CSF NFL | Serum pTau217 (pg/mL) | Clinical symptoms |
| --- | --- | --- | --- | --- | --- | --- | --- | --- |
| IgLON5#1 | M | 59 | MP, PPH | n.a. | 1:1000 (Serum)<br>1:100 (CSF) | n.a. | 0.0722* | sleep apnea, myoclonic jerks, dysarthria, and gait disorder |
| IgLON5#2 | M | 59 | PPH, RTX, Bortezomib | no | 1:1000 (Serum)<br>1:100 (CSF) | 841 | 0.0193 | Gait instability, bulbar symptoms, sleep disturbance, fasciculations, myoclonic jerks, myorhythmia, hyperreflexia |
| IgLON5#3 | F | 35 | IA, RTX | no | 1:3200 (Serum)<br>1:32 (CSF) | 559 | n.a. | Sleep disturbance, fasciculations |
| IgLON5#4 | F | 76 | MP, PPH<br>IVIG | no | 1:1000 (Serum)<br>1:10 (CSF) | 1127 | 0.0246 | Bulbar syndromes, sleep disturbance, respiratory insufficiency, cognitive dysfunction |

### α-IgLON5 AAB detection in patient CSF and serum

Specificity of the purified α-IgLON5 AABs was confirmed by indirect immunofluorescence (IF) performed on HEK293 cells transfected with commercially available IgLON5 expression vector (EUROIMMUN, # 1151-1005-50) and cells with control transfection. α-IgLON5 AAB binding on cells was confirmed at dilutions of 1:100 for CSF and 1:1000 for serum.

### Purification of patient-derived α-IgLON5 AABs

Recombinant human IgLON5-Fc chimera (R&D Systems) was bound to a 1 ml HiTrap® NHS-activated HP column (GE Healthcare) according to manufacturer’s guide. For antibody isolation, eluates of plasmapheresis were loaded overnight onto the column. Then, the column was washed with 15 ml 20 mM Tris (pH 7.5) and 15 ml 0.5 M NaCl in 20 mM Tris (pH 7.5). α-IgLON5 AABs were eluted with 100 mM glycine (pH 2.2) with subsequent neutralization using 1 M Tris (pH 8.8). Finally, the samples were re-buffered into phosphate-buffered saline (PBS) using Vivaspin® 15R ultrafiltration spin columns (Sartorius). For detection of specificity, recombinant IgLON5 or LG1 protein was spotted onto a nitrocellulose membrane (GE Healthcare), membranes were blocked with 1% BSA in PBS, incubated with α-IgLON5 AABs over night at 4°C, washed and incubated with HRP-coupled anti-human IgG. For the isolation of control serum IgGs (pCtrl), 750 μl serum from healthy controls were mixed with 450 μl protein A/G agarose (SantaCruz). IgG fractions were isolated according to manufacturer’s instructions. Purity was checked by Coomassie staining. Protein concentration was determined using BCA protein assay Kit (ThermoFisher Scientific). The non-reactive human monoclonal control AAB (mCtrl; mGO53) was purchased from InVivoBiotech.

### IgG subclass composition of patient-derived AABs

For detection of IgG subclasses, 2 μg of patient-derived α-IgLON5 AABs #1 and pCtrl were dotted on a nitrocellulose membrane (GE Healthcare). Membranes were probed with mouse anti-human IgG1-POD, anti-human IgG2-POD, anti-human IgG3-POD, and IgG4-POD ab (ThermoFisher Scientific), and sheep-anti-human IgG (Seramun). For quantitative determination of subclass distribution across all four isolated α-IgLON5 antibody samples and pCtrl, the Human IgG Subclass ELISA Kit (ThermoFisher Scientific) was used according to the manufacturer’s protocol.

### Relative binding strength quantification using flow cytometry

Binding strength was determined following previously published protocols (*79*). In brief, HEK293T cells were transfected with cDNA plasmids coding for full human IgLON5 protein that co-expressed a myc-tag for transfection efficiency control. Three days post-transfection, cells were harvested and stained with serial dilutions of AABs and c-myc AABs. From live cells with top 30% protein expression (evaluated by c-myc signal), the mean fluorescence intensity (MFI) of Alexa488-coupled goat anti-human IgG (1:500; Life Technologies) was evaluated, and nonlinear regression models [MFI=MFI_max_*IgG conc./ (0.5*MFI_max_+IgG conc.)] under settings for one site specific binding were generated using GraphPad Prism 8 (GraphPad Software Inc.).

### Tissue reactivity screening

Sagittal mouse brain sections were obtained from unfixed tissue, cut on a cryostat (Leica) at 20 μm thickness and mounted on glass slides. Sections were rinsed with PBS, then blocked with PBS containing with 2% bovine serum albumin (BSA) and 5% normal goat serum (NGS) for 1 h at room temperature, and incubated with α-IgLON5#1 and pCtrl overnight at 4°C. After three washes with PBS, goat anti-human IgG-AlexaFluor488 was applied for 2 h at room temperature, followed by staining with DAPI (1:1000 in PBS) for 5 min. Sections were washed with PBS before mounting with Immo-Mount (Epredia) and imaging using an inverted epifluorescence microscope (Leica SPE).

### Epitope mapping in HEK293T cells expressing IgLON5 constructs

To map the epitopes of patient-derived α-IgLON5#1, we cloned human IgLON5 deletion constructs from full-length Myc-DKK-tagged human IgLON5 plasmid (Origene, #225495) using a Q5® Site-Directed Mutagenesis kit (New England Biolabs). Individual Ig domains of IgLON5 (Ig1, Ig2, and Ig3), or combinations thereof (Ig1+Ig2, Ig1+Ig3, and Ig2+Ig3), were expressed in HEK293T cells for 48 h. HEK293T cells were seeded on poly-L-lysine (PLL) coated coverslips and transiently transfected with either full-length IgLON5 or its deletional constructs. After fixation with 4 % paraformaldehyde (PFA) for 10 min at room temperature, blocking with 3% NGS in PBS for 1 h, cells were incubated with α-IgLON5 AABs or pCtrl overnight at 4°C. After washing and incubation with the AlexaFluor488-conjugated anti-human IgG for 1 h at room temperature. Binding of α-IgLON5 AABs was evaluated based on epifluorescence microscope (Leica SPE).

### Intrathecal osmotic pump infusion

Eight weeks-old male C57Bl/6J wild-type mice were randomized for the different treatment groups by an independent investigator. In total, 75 μg (∼3 μg/g b.w.) α-IgLON5#1, IgG from healthy control individual serum (pCtrl), and PBS were delivered continuously into the right lateral ventricle over the course of 14 days, with a flow rate of 0.25 μl/h. The cohort consisted of α-IgLON5 (N = 10), pCtrl (N = 9) and PBS (N = 9) animals. Antibody cerebroventricular infusion was performed unilaterally using osmotic pumps (model 1002, Alzet, Cupertino, CA), which were loaded 24 h prior to surgical implantation. For pump implantation, mice were placed in a stereotaxic frame and a cannula was inserted into the right ventricle (Coordinates: 0.2 mm posterior and ± 1.00 mm lateral from bregma, depth 2.2 mm). The cannula was connected to a pump, which was subcutaneously implanted on the interscapular space of the animals. After surgery, mice were monitored daily to assess symptoms and body weight. Mice were sacrificed after 14 days and brain, and serum were harvested.

### Tissue processing

For protein and RNA analysis, brain and spinal cord tissues were promptly dissected on ice, flash-frozen by placing on dry ice, and stored at −80°C until further use.

For protein extraction, brain and spinal cord tissues were homogenized in ice-cold radioimmunoprecipitation assay (RIPA) buffer (Sigma-Aldrich) containing phosphatase and protease inhibitor cocktail (ThermoFisher Scientific) using a probe homogenizer. After incubating the homogenates on ice for 20 min, samples were centrifuged at 10000xg for 20 min at 4°C. Protein concentration in the supernatant was measured using a BCA protein assay Kit (Pierce). Concentration of each sample was adjusted to 2 μg/μl and then mixed with 6 X SDS-containing Laemmli buffer (ThermoFisher Scientific). Subsequently, they were boiled at 95 °C for 5 min and stored at −20°C until the Western blot analysis.

For Immunohistochemistry (IHC), brains were drop-fixed in 4% PFA for 2 days. For cryoprotection, fixed tissues were sequentially transferred to 10%, 20% and 30% sucrose in PBS with 0.02% sodium azide at 4°C. Tissue was cut into 30 μm thick sections using a cryostat (Leica) and serially collected in 50% glycerol in PBS to be stored at −20°C until used for IHC.

### Western blot

To analyze total and p-Tau levels by Western blot, 10 μg of total protein per sample was loaded onto 4 to 12% bis-tris SDS- polyacrylamide gel (Invitrogen). Following electrophoresis, the separated proteins were transferred to a nitrocellulose membrane. The membrane was blocked with 3% BSA in PBS containing 0.05% of Tween (PBS-T) for 1 h at room temperature. Subsequently, the membranes were incubated with primary antibodies for total Tau, p-Tau and loading control, diluted in the blocking solution, overnight at 4°C. The following day, membranes were washed 3-times for 5 min in PBS-T and then incubated with fluorescent-dye conjugated secondary antibodies, diluted in blocking solution, for 1 h at room temperature. Membranes were then imaged using a Licor imaging system (Odyssey DLx). A full list of antibodies used in Western blot can be found in Table S1.

### Histology

For immunodetection, 30 μm-thick PFA-fixed brain sections were washed free-floating in tris-buffered saline (TBS), permeabilized with 0.3% Triton X-100 in TBS for 20 min at room temperature, and washed 2-times for 5 min in TBS. All incubation steps were performed on an orbital shaker (180 rpm). Sections were blocked in blocking solution (3% NGS in TBS) for 1 h at room temperature. Sections were incubated with primary antibodies diluted in blocking solution at 4°C overnight. Next day, the sections were washed 3-times for 10 min in TBS and incubated with Alexa Fluor conjugated secondary antibody diluted in blocking solution for 2 h at room temperature. Sections were washed 3-times for 10 min in TBS and incubated with DAPI (1:1000 in PBS) for 15 min at room temperature. Sections were then mounted onto microscope slides and covered with coverslip using Fluoroshield™ mounting medium (Sigma-Aldrich). A full list of antibodies used IHC can be found in Table S1. Sections were imaged on a wide field fluorescence microscope (Eclipse-Ti, Nikon) using 10x objective with tiling function. Same imaging parameters were used while imaging the sections across different groups. Mean fluorescence intensity was measured using ImageJ from manually drawn regions of interest (ROIs) in the brain sections (on contralateral site to avoid injection confound). Mean intensity was averaged over 3 - 5 brain sections per animal.

Detection of the spread of infused AABs was done in free-floating brain sections with goat anti-human HRP-conjugated secondary antibody in 3% NGS in TBS for 2 h at room temperature. Sections were washed 3-times for 5 min in TBS and the signal was amplified with a CF® 594 tyramide signal amplification kit (ThermoFisher Scientific).

### Evaluation of serum

Levels of neurofilament light chain (Nfl) was measured from undiluted serum samples using commercial ELISA kit (Cloud-Clone) according to manufacturer’s instructions.

### RNA sequencing

Total RNA was extracted from brain tissue using the miRNeasy Micro Kit (Qiagen) according to the manufacturer’s protocol. RNA integrity was assessed prior to bulk RNA sequencing, which was performed in paired-end mode using the Illumina NovaSeq 6000 platform. Raw FASTQ files were aligned to the GRCm38 reference genome using STAR aligner with default parameters (*80*). The following analysis steps were performed in R (v4.3.0) (R Core Team. R: A language and environment for statistical computing. R Foundation for Statistical Computing) and R Studio (v1.4.1717; RStudio Team. RStudio: Integrated Development for R. RStudio, Inc., Boston, MA (2021)). Genes with fewer than 10 read counts in at least 4 samples were excluded from the analysis, resulting in a filtered dataset of 20,271 genes for downstream processing. Normalization of the count matrix was computed with R/DESeq2 (v1.40.2) (*81*) and a variance stabilizing transformation applied using the DESeq2 vst function at default settings. Differential expression analysis based on the DESeq2 package was performed adjusting p-values according to independent hypothesis weighting from the R/IHW package (v1.28.0) (*82*) and applying apeglm shrinkage from the R/apeglm package (v1.22.1) (*83*). DEGs were defined based on a fold change threshold > 2 and a p-value threshold of < 0.05. A gene set enrichment analysis (GSEA) was performed with the transformed data as the input using R/fgsea (v1.26.0) (*84*), whereby the Gene Ontology (GO) (*85*, *86*), and the Molecular Signature Database (MSigDB) (*87*, *88*), Hallmark gene set was used.

### Preparation of mouse primary neurons

Primary neurons were prepared from hippocampi dissected from P0-P1 wildtype mice of either sex. Hippocampi were dissected in ice-cold HBSS (Millipore) containing 1% P/S. Then, hippocampal tissue was digested in enzyme solution containing DMEM (ThermoFisher Scientific), 3.3 mM cysteine, 2 mM CaCl2, 1 mM EDTA, and papain (20 U/ml, Worthington) at 37°C for 30 min. Papain reaction was inhibited by incubating digested hippocampal tissue in DMEM containing 10% FBS (ThermoFisher Scientific), 1% P/S, 38 mM BSA, and 95 mM trypsin inhibitor at 37°C for 5 min. Cells then were triturated in complete Neurobasal-A (NBA) medium containing 10% FBS, 2% B-27, 1% Glutamax, 1% penicillin/streptomycin (P/S) and seeded in PLL (0.1 mg/mL) coated glass bottom μ-slide 8-well imaging dishes (ibidi) at a seeding density of ∼30000 cells/cm^2^ for IF experiments. For calcium (Ca^2+^) imaging, neurons were seeded in 96-well clear bottom black microplates (Corning) at a seeding density of 125000 cells/cm^2^ and maintained at 37°C and 5% CO_2_ until used for experiments. After 3 h seeding the neurons, the media was completely changed to FBS and phenol-red free complete NBA medium. Every second day, one-fifth of the media was replaced by fresh complete NBA medium.

### Differentiation of human neurons

Human neuronal stem cells (hNSCs) were differentiated using an inducible Neurog2 hNSCs through doxycycline induction. hNSCs were received from BIH Core Unit pluripotent Stem Cells and Organoids (CUSCO), Charité-Berlin. Cells were seeded in laminin 521 (Biolamina)-coated 8-well plates. The composition of the differentiation medium was: NBA, Neural Induction Supplement, Advanced DMEM/F12 (Gibco), supplemented by antibiotic (Gibco) and doxycycline (Sigma-Aldrich). On Day 4, the medium change was performed by the medium supplemented with AraC (Sigma-Aldrich), to reduce the number of non-neuronal cells. The differentiation medium was refreshed every day, the cells were cultivated for 43 days in 37°C and 5% CO₂.

### Antibody binding curves on the neuronal surface

Primary mouse hippocampal neurons (DIV12) were fixed with 4% PFA for 15 min at room temperature and blocked with 3% NGS in PBS for 1 h. For surface staining with AABs, the permeabilization step was omitted. Neurons were then incubated overnight at 4°C with increasing concentrations (0.01, 0.1, 1, 10 μg/ml) of α-IgLON5 AABs or pCtrl diluted in blocking buffer. The next day, cells were washed with PBS and incubated for 2 h at room temperature with Alexa Fluor 488-conjugated α-human secondary antibodies. After additional PBS washes, neurons were permeabilized with 0.3% Triton X-100 in PBS for 20 min, re-blocked, and stained overnight at 4°C with MAP2 primary antibodies. Alexa Fluor 555-conjugated secondary antibodies were applied for 2 h at room temperature the following day. Nuclear staining was performed using DAPI (1:1000 in PBS, 10 min at room temperature), followed by a final PBS wash. Neurons were imaged using a spinning disk confocal microscope (Nikon CSU-X) equipped with a 10X objective lens. Quantitative analysis of antibody binding was done by measuring the MFI within dendritic regions. Dendrites were identified based on MAP2-positive staining and delineated using CellProfiler software. The MFI was measured for each treatment concentration using ImageJ software. Concentration-dependent binding curves were then generated.

### Live staining (surface antigen binding)

Primary mouse hippocampal neurons (DIV12) were treated with 1 μg/ml α-IgLON5 AABs or control antibodies and kept at 37°C and 5% CO_2_ for 60 min. Then, neurons were washed with PBS before fixation with 4% PFA in PBS for 15 min, and then washed with TBS for 10 min at room temperature. After fixation, neurons were blocked with 3% NGS for 1 h at room temperature and incubated with anti-human 488 secondary antibodies for 2 h at room temperature. After 3-times 10 min PBS washes, nuclei were stained with DAPI (1:1000 in PBS) for 10 min. Dishes were imaged with laser scanning confocal microscope (Nikon, A1Rsi+) with a x60 oil objective.

Co-localization with synaptic markers: After the surface antigen labelling as described above, neurons were permeabilized with PBS containing 0.3% Triton X-100 for 20 min at room temperature. After two consecutive washing steps with excess PBS, neurons were blocked again with 3% NGS for 1 h. Neurons then were incubated with primary antibodies, diluted in blocking solution, for microtubule (MAP2) and pre- (Synapsin1), or post- (PSD95) synaptic markers overnight at 4°C. Secondary antibody and DAPI stainings were performed as described above. Dishes were imaged with a laser scanning confocal microscope (Nikon, A1Rsi+) with a x60 oil objective using the z- stack function (z-step: 0,5 μm). Same imaging settings were used between each biological replicate (N=3) to allow the comparison for co-localization analysis, which was performed using standard procedures and custom-made macros in ImageJ.

### IgLON5 knock-down in Neuro2a cells and primary mouse neurons

To knock down mouse IgLON5, pAAV-U6-shRNA vectors encoding either IgLON5-shRNA or scramble-shRNA, both expressing a BFP marker, were obtained from VectorBuilder. Knock-down efficiency was first validated in Neuro2a mouse neuroblastoma cells (ATCC CCL-131). Cells were cultured to ∼60% confluence in DMEM supplemented with 10% FBS, 1% Pen/Strep, 2 mM L-glutamine, and 1% NEAA, and transfected using Lipofectamine 2000 per the manufacturer’s protocol. After 24 h, cells were fixed with 4% PFA and surface IgLON5 staining was performed using α-IgLON5#1. Cell membranes were labeled with CellBrite-Red (Biotium). Imaging was conducted on a spinning disk confocal microscope (Nikon CSU-X, 40X objective). Following validation, AAV2/9 viral particles were produced by the Viral Core Facility at Charité (IgLON5-shRNA: 8.26×10¹² VG/ml; scramble-shRNA: 9.86×10¹² VG/ml). At DIV5, primary mouse hippocampal neurons neurons were transduced with serial dilutions (0.001-0.5 μl) of IgLON5-shRNA or scramble-shRNA AAVs. At DIV12, cells were fixed and stained for surface IgLON5 using α-IgLON5#1, followed by MAP2 staining for dendritic visualization. Imaging was performed on a spinning disk confocal microscope (Nikon CSU-X, 10X objective). Quantification of α-IgLON5 AAB binding was done by measuring MFI in dendritic regions of IgLON5-shRNA and control neurons.

### Tau missorting analysis in neurons

Primary neurons were treated with different concentrations of α-IgLON5 AABs or control antibodies for either 2 days or 5 days. For Ca^2+^ depletion experiments, neurons were treated with 10 μM cell permeable Ca^2+^ chelator, EGTA-AM, (AAT Bioquest) 20 min prior to the antibody treatment. After the treatments, neurons were fixed at DIV9-12 with 4% PFA for 15 min and immunolabeled for total Tau and MAP2 by following the IF protocol described above. Antibody specifications are listed in Table S1. Neurons were imaged with a spinning disc confocal microscope (Nikon, CSU-X), using a 40X oil objective. Identical imaging settings were used for all conditions. ROIs were manually defined for the soma and nucleus based on signals from the MAP2 and DAPI channels, respectively. Mislocalization of Tau into the somatodendritic compartment was quantified by calculating MFI from the Tau channel using the following equation:

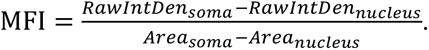

### FTD-Tau aggregation analysis in neurons

Primary neurons (DIV5) were transduced with AAV2/9 serotype viral particles encoding GFP-tagged human full-length Tau (2N4R isoform) carrying two FTD-associated mutations (ΔK280. P301L, P301L/S320F) under hSyn1 or CAG promoter. At DIV12, neurons were treated with either pCtrl or α-IgLON5 AABs (1 μg/ml) for 2 days, fixed on DIV14, and immunolabeled for MAP2 and DAPI. GFP-TauΔK280 and GFP-TauP301L expressing neurons were treated with AABs for 7 days. Tau aggregates were manually counted and the percentage of neurons with tangle-like Tau aggregates calculated as # tangles/ # DAPI+ nuclei.

### Cytotoxicity assay (Lactose dehydrogenase, LDH)

To assess cytotoxicity, neuronal culture supernatants were collected, centrifuged at 1000 g for 5 min, and analyzed using the CyQUANT LDH assay (ThermoFisher Scientific) per the manufacturer’s instructions. Absorbance was measured at λ490 nm in Tecan Infinite® M Plex plate reader.

### Ca^2+^ imaging in cultured neurons

For Ca^2+^ imaging in cultured hippocampal neurons, AAV2/9 serotype was used for neuron-specific expression of GCamp6f under the human synapsin (hSyn) promoter. AAVs expressing hSyn:mRuby2-P2A-GCamp6f were produced by the Viral Core Facility of Charité Universitätsmedizin, Berlin (Catalog number: BA-026e). AAVs (0.16 μl of 1,04×10^13^ VGs/ml) were added to hippocampal neurons cultured in 96-well plates at DIV5. Ca^2+^ recordings were performed at DIV13-15. Additionally, immunostaining for c-FOS, an immediate early gene marker of neuronal activation, was performed in treated neurons to confirm AAB-induced hyperactivity. Ca^2+^ transient recordings: Neurons in 96-well plates expressing GCamp6f were imaged using a wide field fluorescence microscope (Eclipse-Ti, Nikon) with a 10x objective. Imaging dishes were placed in the live-cell imaging chamber of the microscope to maintain an environment of 37°C, 5% CO_2_ and ∼95% humidity. α-IgLON5 AABs or control antibodies were added to the wells by direct application. Bicuculline (30 μM for 5 minutes) and TTX (0.5 μM for 10 minutes) were included in the recordings as controls. Ca^2+^ imaging was done in the green channel and mRuby expressing neurons were visualized in the red channel. Ca^2+^ transients were recorded for 180 sec at ∼8 Hz. Two to three wells in three independent cultures were analyzed per condition.

<u>Analysis:</u> ROIs were created for each soma in the red channel using a custom written script in Cell Profiler. Created ROIs were placed over each frame of the video recorded in the green channel and mean intensity over time was measured using ImageJ. The .csv files containing mean fluorescent intensity from each image of the time-lapse videos were processed using the FluoroSNAP application for ΔF/F0 conversion and spike detection (*89*). In short, .csv files were uploaded to FluoroSNAP and the function “Convert raw fluorescence data to ΔF/F0” was used to compute baseline fluorescence by taking the average of the 50th percentile of the signal across a 60 s time window. Ca^2+^ transients from individual ROIs were detected with a template-based approach. This method identifies events by comparing a moving window of the Ca^2+^ signal to a predefined library of Ca^2+^ waveform templates. Events were detected with a similarity threshold of 0.75 and a minimum amplitude of 0.01. Hyperactivity criteria was calculated for each time point independently by taking the mean±SD of the pCtrl group as a reference value for normal activity: Spike frequencies larger than [mean + 2*SD] of pCtrl were considered as hyperactivity (*90*, *91*).

### Electrophysiology in primary autaptic cultures

Isolated primary neurons on micro islands of glial cells (“autaptic cultures”) were prepared from wildtype mice as recently described (*92*), with slight modifications. Briefly, 300 µm spots of a growth-permissive substrate mix of 0.7 mg/ml collagen and 0.1 mg/ml poly-D-lysine were printed on glass coverslips coated with agarose. Astrocytes were seeded onto these coverslips in DMEM (ThermoFisher Scientific), supplemented with 10% fetal calf serum and 0.2% P/S (Invitrogen). After formation of glia micro-islands, DMEM was replaced with NBA supplemented with 2% B27 and 0.2% P/S. Hippocampal neurons prepared from P0 mice were added at a density of 370 cells/cm^2^. Prior to electrophysiological recordings, autaptic cultures were treated with antibodies at a final concentration of 0.1 µg/ml for 3 days.

Electrophysiological recordings: Neurons were recorded at DIV13-17 at room temperature on an IX73 inverted microscope (Olympus) using a Multiclamp 700B amplifier under the control of a Digidata 1550 AD board and Clampex 10 software (all Molecular Devices). Data were acquired at 10 kHz and filtered at 3 kHz, and series resistance was compensated at 70%. The extracellular solution contained 140 mM NaCl, 2.4 mM KCl, 10 mM HEPES, 10 mM glucose, 2 mM CaCl_2_, and 4 mM MgCl_2_ (pH adjusted to 7.3 with NaOH, 300 mOsm). The intracellular solution contained 136 mM KCl, 17.8 mM HEPES, 1 mM EGTA, 4.6 mM MgCl_2_, 4 mM Na_2_ATP, 0.3 mM NaGTP, 12 mM disodium phosphocreatine, 50 U/ml creatine phosphokinase, pH adjusted to 7.3 with KOH, 300 mOsm. Autaptic neurons were recorded in whole-cell voltage clamp mode using thick-walled borosilicate pipettes with a tip resistance of 3-4 MΩ. Membrane potential was set to −70 mV. Paired EPSCs were evoked every 5 s by triggering two unclamped action potentials with 40 ms interstimulus interval using 1 ms depolarizations of the soma to 0 mV. The readily-releasable pool of synaptic vesicles was determined by application of a hypertonic 500 mM sucrose solution for 10 sec. Electrophysiological recordings were analyzed using AxoGraph. The vesicular release probability was calculated as the ration of the charge of the sucrose evoked response by the average charge of six EPSCs prior to the sucrose application. The paired-pulse ratio was calculated as the ratio from the second and first EPSC amplitude.

### Antibody internalization experiments

α-IgLON5 AABs and pCtrl were conjugated with endosomal-pH sensitive red-fluorescent dye (pHrodo^TM^ iFL Red STP ester, ThermoFisher Scientific) following the manufacturer’s instructions. pHrodo^TM^-conjugated antibodies were applied at 5 μg/ml to primary mouse hippocampal neurons (DIV12) for 2 days at 37°C. Images were captured on a spinning disc confocal microscope (Nikon, SoRa CSU-W1).

### Cell surface clustering of α-IgLON5 AABs

Live primary mouse neurons (DIV12) were treated with α-IgLON5#1 or pCtrl (1 μg/ml) or left untreated (NT), and incubated at 37°C, 5% CO₂ for 60 min. After incubation, cells were washed with PBS, fixed with 4% PFA, and surface-stained using α-IgLON5 AABs as described under “Antibody Binding Curves on the Neuronal Surface.” MAP2 staining was used to visualize dendrites. Imaging was performed on a Nikon A1Rsi+ laser scanning confocal microscope using a 60X oil objective with z-stack (0.5 μm steps), applying consistent settings across biological replicates. IgLON5 cluster density, size and radius were quantified using a custom CellProfiler pipeline.

### Fab fragment preparation and cell treatments

Fab fragments were generated from α-IgLON5#1 using a commercially available Fab preparation kit (ThermoFisher Scientific) following the manufacturer’s protocol. Briefly, IgG was digested with papain to cleave the Fc region and isolate antigen-binding Fab fragments. The resulting fragments were separated via SDS-PAGE to confirm digestion efficiency. For all Fab assays, neurons were treated with equimolecular concentrations of α-IgLON5#1, α-IgLON5#1 fab fragment or pCtrl.

### Antibody-mediated cell surface proximity biotinylation

Primary mouse neurons (DIV12) were treated with α-IgLON5#1 or pCtrl (1 μg/ml) for 60 min at 37°C. After washing with PBS, cells were fixed with 4% PFA for 10 min at room temperature, quenched with 100 mM glycine (10 min), and treated with 0.5% H₂O₂ (10 min) to block endogenous peroxidase. Following PBS washes, cells were blocked with 5% biotin-free BSA for 45 min at room temperature, then incubated with HRP-conjugated α-human secondary antibodies for 1 h. Surface biotinylation was performed using 250 μM Biotin-Tyramide (Iris Biotech) and 50 μM H₂O₂ for 1 min at t room temperature, followed by quenching with Trolox and sodium L-ascorbate. Cells were washed and prepared for imaging.

<u>Immunofluorescence assessment of biotinylation:</u> Cells were stained with Alexa Fluor 555-streptavidin to detect biotinylated proteins and Alexa Fluor 488-conjugated α-human antibodies to visualize bound α-IgLON5 AABs. Nuclei were counterstained with DAPI. Imaging was conducted on a spinning disk confocal microscope (Nikon CSU-X, 40X oil objective), and co-localization analysis was performed using intensity profiling in ImageJ.

<u>Streptavidin pulldown and mass spectrometry:</u> Neurons were seeded into 6-well plates (500,000 cells/well). After surface biotinylation, cells were scraped with PBS, with three wells (∼1.5 million cells) pooled per condition. Pooled cells were pelleted by centrifugation at 8000×g for 5 minutes at room temperature to remove supernatant. Cell pellet was lysed in ice cold RIPA buffer (with 0.5% SDS, protease/phosphatase inhibitors). Lysates were sonicated, incubated on ice for 20 min, and boiled (99°C, 1 h) for de-crosslinking. Soluble proteins were collected by centrifugation (14,000×g, 20 min, 4°C), and protein concentration was determined using a BCA assay (ThermoFisher Scientific). For pulldown, streptavidin magnetic beads (BioLabs)were equilibrated in RIPA buffer. For each pulldown, 180 μg of protein lysate was incubated with beads overnight at 4°C. Next day, beads were washed with ice cold RIPA buffer and biotinylated proteins were separated from the beads through competitive elution by adding 10 mM Biotin-Tyramide to the bead slurry and by boiling at 95°C for 10 minutes. The eluted proteins were stored at −80°C for subsequent mass spectrometry analysis. In addition to the pulldown samples, 30 μg of total protein lysates from α-IgLON5#1 or pCtrl treated neurons were included in the mass spectrometry analysis to ensure that the protein composition in the input samples was comparable across treatment conditions.

<u>Mass spectrometry sample preparation and analysis:</u> Pulldown (∼110 μl) and total lysate (∼50 μl) samples were thawed on ice and treated with 0.2 μl and 0.5 μl nuclease, respectively. Samples were diluted 1:1 with 50 mM ammonium bicarbonate (ABC). SP3-beads (Sera-Mag SpeedBeads) were prepared as described before (*93*), and 10 μl of beads per pulldown and 15 μl of beads per lysates samples were added. Acetonitrile was added to a final concentration of 70% (v/v), and samples were incubated for 30 min at 24°C with shaking (1000 rpm), and supernatants were discarded. Beads were resuspended in 20 μl 50 mM ABC and reduced with 3 μl 200 mM DTT at 45°C for 20 min (1200 rpm). After cooling to room temperature, 7 μl of 400 mM iodoacetamide was added, and the samples were incubated at 24°C with shaking at 1200 rpm for 30 minutes, followed by addition of 3 μl of 200 mM DTT. Freshly prepared 10 μl SP3-beads were added to the sample still containing previous beads and ACN was added to a final concentration of 70% (v/v). After incubation for 30 minutes shaking at 24 °C, the beads were washed 4-times with 80% ethanol, and the supernatant was discarded. For digestion, 20 μl of digestion mix (0.03 μg/μl trypsin, 0.015 μg/μl Lys-C, 50 mM ABC) was added per 20 μg of protein. The samples were gently spun and incubated overnight at 37°C. Supernatants were subsequently filtered thorough pre-equilibrated (0.1% formic acid (FA)) Spin-X 0.22 μm filters and collected in a fresh tube. The remaining beads were resuspended in 20 μl of 0.1% FA, sonicated twice for 30 seconds in a water bath, and eluates were filtered through Spin-X 0.22 μm filters. Both eluates were combined and dried by vacuum centrifugation. Dried peptides were reconstituted in 13 μl of 0.1% FA. In total, 6 μl of pulldowns and 350 ng of peptides per lysate were injected via a nanoElute nanoHPLC system (Bruker, Germany) coupled to a TimsTOF pro mass spectrometer (Bruker, Germany) with a CaptiveSpray ion source (Bruker, Germany). Samples were separated on an in-house packed C18 analytical column (15 cm × 75 μm ID, ReproSil-Pur 120 C18-AQ, 1.9 μm, Dr. Maisch GmbH) using a 300 nl/min gradient of water and ACN (B) containing 0.1% FA (0 min, 2% B; 2 min, 5% B; 62 min, 24% B; 72 min, 35% B; 75 min, 60% B) at a column temperature of 50°C. Spectra were acquired with Data Independent Acquisition Parallel Accumulation–Serial Fragmentation (diaPASEF). Ion accumulation and separation using Trapped Ion Mobility Spectrometry (TIMS) was set to a ramp time of 100 ms. One scan cycle consisted of a TIMS full MS scan and 26 windows with a width of 27 m/z covering a m/z range of 350-1002 m/z. Two windows were recorded per PASEF scan, resulting in a cycle time of 1.4 s.

<u>Mass spectrometry data processing:</u> MS data were analysed with DIA-NN version 1.8.2 (*94*). A library free search was performed against a mouse FASTA database including common contaminants. Methionine oxidation and acetylation of protein N-termini were set as variable modifications, carbamidomethylation of cysteine residues as fixed modification. The match between runs-option was enabled while data normalization was disabled. Charge states from two to four with a m/z range of 300 to 1400 were considered. Trypsin with up to two missed cleavages was set as digestion condition. Mass accuracy and ion mobility settings were set to automatic. Statistical data processing was performed with the freeware tool Perseus version 2.0.11 (*95*).

### Statistical analysis

All data plotting and statistical analyses were performed in GraphPad Prism 10. Comparisons between two groups were performed using the unpaired t-tests, while One-way ANOVA followed by Tukey’s test used for multiple group comparisons, as specified in the figure legends. Statistical significance was denoted as follows: ****p<0.0001, ***p<0.001, **p<0.01, *p<0.05.

### Data and code

RNA seq data generated in this study are available in the Gene Expression Omnibus (GEO) at https://www.ncbi.nlm.nih.gov/geo/query/acc.cgi?acc=GSE298951, under the accession number <u>GSE298951</u>, with the token: <u>mduriemyzlsvdej</u>. Lists of differentially expressed gene can be found in Data S1. Code for RNA seq analysis is available at https://gitlab.dzne.de/grundschoettelp/iglon5-autoimmune-antibody-influence-on-neurons.

The proteomics data have been deposited to the ProteomeXchange Consortium via the PRIDE partner repository with the dataset identifier PXD066225 (token i5YAGUXZ5Yv4, username, password: rbRRDufu6h6K). Lists of differentially enriched proteins can be found in Data S3.

## Acknowledgments

We thank Britta Eickholt for providing us with access to instruments and equipment of the Charité Biochemistry Department, the Advanced Medical Bioimaging Core Facility (AMBIO) of the Charité, which enabled us to perform most microscopy on brain sections and neuron/cell cultures, the Viral Core Facility (VCF) of the Charité for virus production, and the PRECISE sequencing platform for RNA sequencing of mouse brain tissue.

## Author contributions

Conceptualization: SW, HP, PK

Methodology: BA, CK, AG, DR, CCG, SLLD, ADB, AN, PG, TU, EW, JL, JW, JN, VT, JP, PT, BRR, KN, SFL, SH, ES

Visualization: BA, SW, CK Funding acquisition: SW, HP, BRR Project administration: SW, HP Writing – original draft: BA, SW

Writing – review & editing: All authors

**Diversity, equity, ethics, and inclusion:** All authors state their clear commitment to promote and support diversity, inclusion, equality, and equity in the clinical and research community.

## Data and materials availability

All data are available in the main text or the supplementary materials.

## Supplementary Materials

Supplementary Text Figs. S1 to S5

Table S1

References (*75–77*)

Data S1 to S4

