## Supplementary Material, Text + Figures S1-S6 + Table S1 for "IgLON5 autoimmune antibodies activate Tau via neuronal hyperactivity"

**Supplementary Text**

Our data suggests that α-IgLON5 AABs induce pathogenic effects in the brain through neuroinflammation, neuronal physiology, and Tau. This combination of insults likely determines disease progression and clinical manifestation of individual anti-IgLON5 disease patients (*22*, *75*, *76*). We find that all polyclonal α-IgLON5 AAB pools have a similar IgG-subclass composition and similar amounts of IgM. The subclass composition of total plasma IgGs of these patients could still be similar to previous reports though (mostly subclasses IgG1 and IgG4) (*19*, *22*). Notably, the IgG subclass composition of α-IgLON5 AABs can have large effects on neuroinflammation and related neurotoxicity, and therefore could also influence Tau pathology in anti-IgLON5 disease patient brain. For the acute effects of α-IgLON5 AABs on neuronal activity and Tau missorting, detected herein in cultured neurons, the influence of IgG subclass composition may be of less importance.

Parenchyma concentration and distribution of cerebroventricularly delivered antibodies in our mouse model are different from that in anti-IgLON5 disease patients. Hence, the brain regions affected by the increase in phosphorylated Tau differ between the animal model and human brains. For example, we observe Tau phosphorylation selectively in the hippocampus, a region of high α-IgLON5 AAB concentration in our model. In these mice, accumulation of Tau-pS396/pS404 occurred in somata of DG granule cells and their projections onto CA3, important structures for learning and memory, and in commissural fibres in corpus callosum, output structures of CA3. Similar hippocampal connections were recently suggested to be functionally affected by intraventricularly delivered α-IgLON5 AABs (*51*, *52*), supporting our observations, and are also affected by Tau in AD (*77*, *78*). Notably, upon two weeks of AAB delivery, we did not detect neurotoxicity or Tau aggregation in wildtype mice. We suspect that long-term (e.g., 1-6 months) or repeated cerebroventricular infusion of non-toxic AAB doses may be needed to induce full Tau pathology and neurodegeneration.

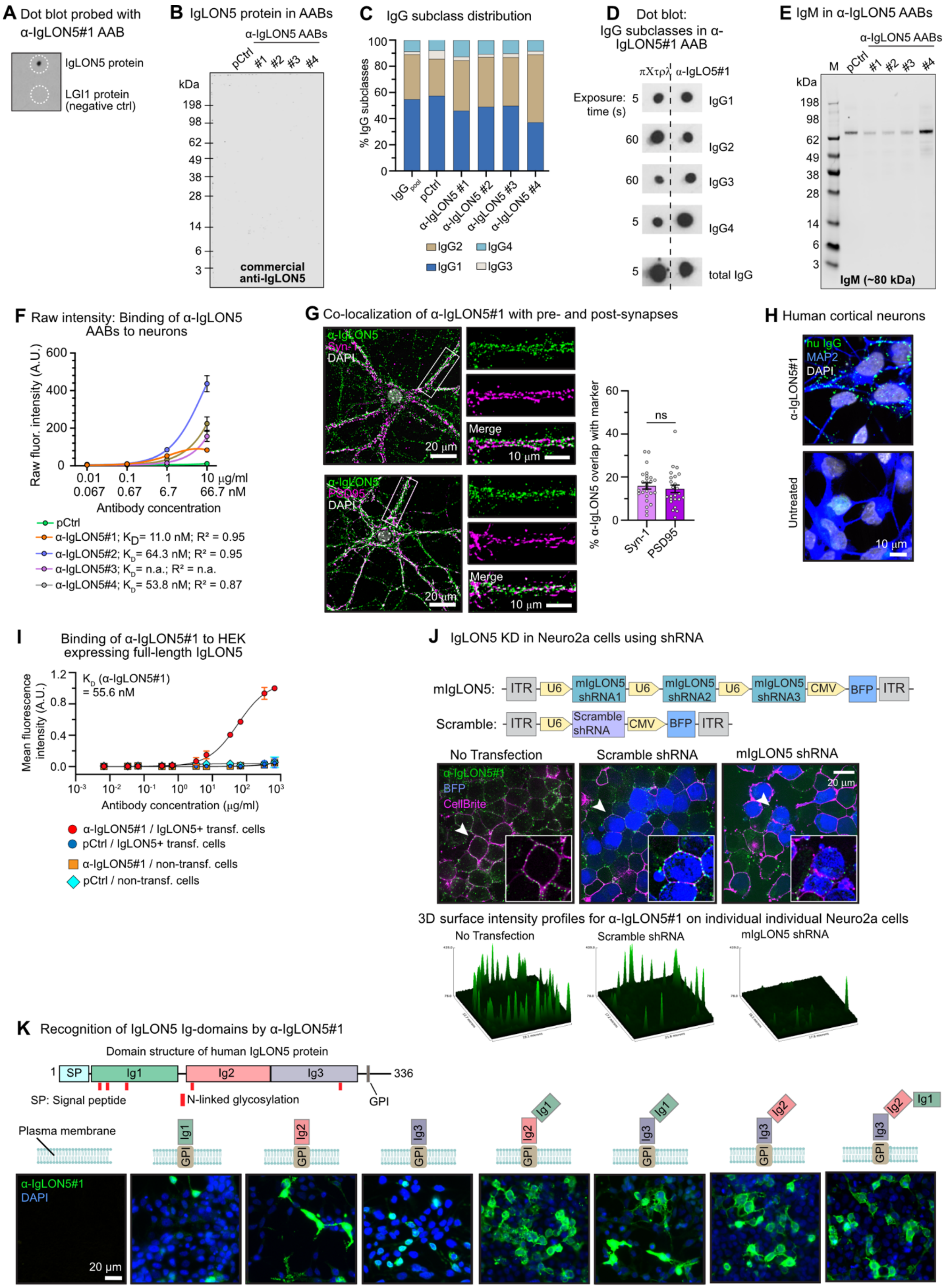
A Dot blot probed with α-IgLON5#1 AAB
IgLON5 protein
LGI1 protein (negative ctrl)
B IgLON5 protein in AABs
α-IgLON5 AABs
pCtrl #1 #2 #3 #4
kDa
198
98
62
49
38
28
14
6
3
commercial anti-IgLON5
C IgG subclass distribution
% IgG subclasses
100
80
60
40
20
0
IgG pool
pCtrl
α-IgLON5 #1
α-IgLON5 #2
α-IgLON5 #3
α-IgLON5 #4
IgG2 IgG4
IgG1 IgG3
D Dot blot: IgG subclasses in α-IgLON5#1 AAB
πΧτρλ α-IgLO5#1
Exposure: time (s)
5 IgG1
60 IgG2
60 IgG3
5 IgG4
5 total IgG
E IgM in α-IgLON5 AABs
α-IgLON5 AABs
M pCtrl #1 #2 #3 #4
kDa
198
98
62
49
38
28
14
6
3
IgM (~80 kDa)
F Raw intensity: Binding of α-IgLON5 AABs to neurons
Raw fluor. intensity (A.U.)
600
400
200
0
0.01 0.1 1 10 µg/ml
0.067 0.67 6.7 66.7 nM
Antibody concentration
pCtrl
α-IgLON5#1; KD= 11.0 nM; R² = 0.95
α-IgLON5#2; KD= 64.3 nM; R² = 0.95
α-IgLON5#3; KD= n.a.; R² = n.a.
α-IgLON5#4; KD= 53.8 nM; R² = 0.87
G Co-localization of α-IgLON5#1 with pre- and post-synapses
α-IgLON5
Syn-1
DAPI
Merge
20 µm
10 µm
α-IgLON5
PSD95
DAPI
Merge
20 µm
10 µm
% α-IgLON5 overlap with marker
60
40
20
0
ns
Syn-1
PSD95
H Human cortical neurons
α-IgLON5#1
hu IgG
MAP2
DAPI
Untreated
10 µm
I Binding of α-IgLON5#1 to HEK expressing full-length IgLON5
Mean fluorescence intensity (A.U.)
1.2
0.8
0.4
0.0
KD (α-IgLON5#1) = 55.6 nM
10⁻³ 10⁻² 10⁻¹ 10⁰ 10¹ 10² 10³
Antibody concentration (µg/ml)
α-IgLON5#1 / IgLON5+ transf. cells
pCtrl / IgLON5+ transf. cells
α-IgLON5#1 / non-transf. cells
pCtrl / non-transf. cells
J IgLON5 KD in Neuro2a cells using shRNA
mIgLON5: ITR U6 mIgLON5 shRNA1 U6 mIgLON5 shRNA2 U6 mIgLON5 shRNA3 CMV BFP ITR
Scramble: ITR U6 Scramble shRNA CMV BFP ITR
No Transfection
Scramble shRNA
mIgLON5 shRNA
α-IgLON5#1
BFP
CellBrite
20 µm
3D surface intensity profiles for α-IgLON5#1 on individual individual Neuro2a cells
No Transfection
Scramble shRNA
mIgLON5 shRNA
K Recognition of IgLON5 Ig-domains by α-IgLON5#1
Domain structure of human IgLON5 protein
1 SP Ig1 Ig2 Ig3 336
SP: Signal peptide
N-linked glycosylation
GPI
Plasma membrane
Ig1 Ig2 Ig3
GPI
α-IgLON5#1
DAPI
20 µm

**Fig. S1. Supplementary Figure S1. α-IgLON5 AABs characterization.**

(**A**) Dot blot of recombinant IgLON5 and LGI (negative ctrl) protein shows specific reactivity of AAB α-IgLON5#1 against IgLON5 protein. (**B**) Western blot, probed with commercial anti-IgLON5 antibody, shows no IgLON5 protein in AAB preparations. (**C**) ELISA-based quantification of IgG subclasses in α-IgLON5 AAB and pCtrl preparations and, for comparison, in a commercial human serum IgG pool ($IgG_{pool}$). (**D**) Dot blot for IgG subclasses ($IgG_1$, $IgG_2$, $IgG_3$ and $IgG_4$) in α-IgLON5#1 and pCtrl. Note, different exposure times for IgG subclass secondaries were used. (**E**) Western blot showing IgM levels in AAB preparations. (**F**) Raw fluorescence intensity values for anti-human-Alexa488 detecting IgLON5 AAB binding to neuronal cultures. Note, values differ between curves because binding curves were determined at different experimental days using different microscope. (**G**) Partial co-localization of α-IgLON5#1 with pre-synaptic (Synapsin-1) and post-synaptic (PSD-95) markers. Scale bars as indicated. Data points are individual images analyzed. N = 3 independent experiments with 3-5 technical repeats each. Student t-test, ns = not significant. (**H**) Binding of α-IgLON5#1 to human iPSC-derived neurons. (**I**) Binding affinity of α-IgLON5#1 to HEK cells recombinantly expressing full-length human IgLON5 (IgLON5+; $EC_{50}$ = 8 μg/ml; $K_D$ = 55.6 nM) and of controls. Fluorescence quantified by FACS of cells after treatment. Data shown as mean±SEM, N = 3 independent experiments. (**J**) shRNA-mediated IgLON5 knockdown in Neuro2a cells. Reduced α-IgLON5#1 immunofluorescence indicates knockdown in mIgLON5. Cell membranes were stained with CellBrite. (**K**) Binding of α-IgLON5#1 to different IgLON5 IgG domains. Domain structure of human IgLON5 protein (336 amino acids) is shown, consisting of a secretion signal peptide (SP), three extracellular Ig domains (Ig1/2/3), and a C-terminal GPI-anchor. Example images show binding of α-IgLON5#1/ anti-human IgG-Alexa488 to HEK cells recombinantly expressing full-length IgLON5 protein or (combinations of) individual Ig domains. Scale bars as indicated.

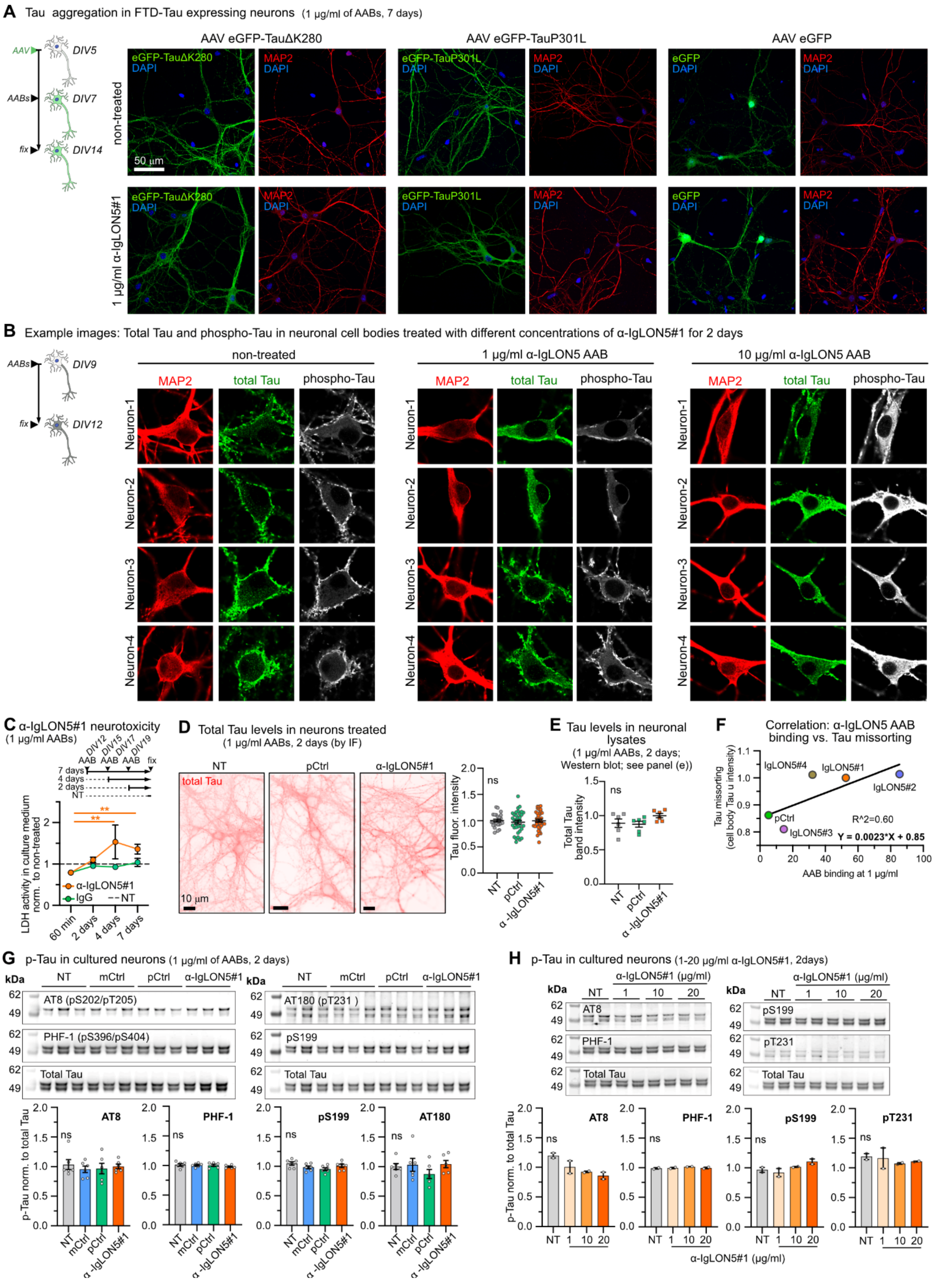

A Tau aggregation in FTD-Tau expressing neurons (1 µg/ml of AABs, 7 days)
AAV eGFP-TauΔK280
AAV eGFP-TauP301L
AAV eGFP
non-treated
1 µg/ml α-IgLON5#1
50 µm
B Example images: Total Tau and phospho-Tau in neuronal cell bodies treated with different concentrations of α-IgLON5#1 for 2 days
non-treated
1 µg/ml α-IgLON5 AAB
10 µg/ml α-IgLON5 AAB
MAP2
total Tau
phospho-Tau
C α-IgLON5#1 neurotoxicity (1 µg/ml AABs)
LDH activity in culture medium norm. to non-treated
D Total Tau levels in neurons treated (1 µg/ml AABs, 2 days (by IF)
Tau fluor. intensity
E Tau levels in neuronal lysates (1 µg/ml AABs, 2 days; Western blot; see panel (e))
Total Tau band intensity
F Correlation: α-IgLON5 AAB binding vs. Tau missorting
R^2=0.60
Y = 0.0023*X + 0.85
AAB binding at 1 µg/ml
G p-Tau in cultured neurons (1 µg/ml of AABs, 2 days)
H p-Tau in cultured neurons (1-20 µg/ml α-IgLON5#1, 2days)
p-Tau norm. to total Tau
α-IgLON5#1 (µg/ml)

**Fig. S2. Tau in AAB-treated neurons.**

(**A**) Representative images of neurons expressing single FTD-mutant Tau variants, eGFP-TauDK280 and eGFP-TauP301L, or eGFP alone treated with α-IgLON5#1 or not. In no condition, Tau aggregates could be found. (**B**) Representative images of neuronal cell bodies stained for MAP2, total Tau and phospho-Tau after treatment with 0, 1, or 10 µg/ml α-IgLON5#1 for 2 days. (**C**) Time-dependent toxicity (LDH assay) in neurons treated with 1 µg/ml α-IgLON5#1 for 60 min, 2 days, 4 days, or 7 days. N = 3 experiments with 3-4 replicates. Two-way ANOVA with Sidak post-test. (**D,E**) Total Tau levels in neurons upon treatment with α-IgLON5#1 or pCtrl (1µg/ml) for 2 days, measured by immunofluorescence (D) and Western blot (E). N = 3 experiments with 3 replicates. One-way ANOVA with Tukey post-test. (**F**) Correlation (linear regression for $R^2$ and p-values) between α-IgLON5 AAB binding at 1 µg/ml and missorted Tau in cell bodies after treatment with 1 µg/ml AABs for 2 days. Data points represent the different patient-derived α-IgLON5 AABs. (**G**) Western blots for phospho-Tau (p-Tau pS202/pT205 (AT8), pS396/pS404, pS199, and pT231 (AT180)) and total Tau levels in lysates from neurons treated with 1 µg/ml α-IgLON5#1 or control antibodies for 2 days. mCtrl is a non-reactive human monoclonal control antibody (mGO). N = 2 experiments with 3 technical replicates per condition. One-way ANOVA with Tukey post-test. (**H**) Western blots for phospho-Tau (p-Tau pS202/pT205 (AT8), pS396/pS404, pS199, and pT231 (AT180)) and total Tau levels in lysates from neurons treated with 1, 10, or 20 µg/ml α-IgLON5#1 or control antibodies for 2 days. N = 2 independent experiment with 3 replicates per condition. One-way ANOVA with Tukey post-test.

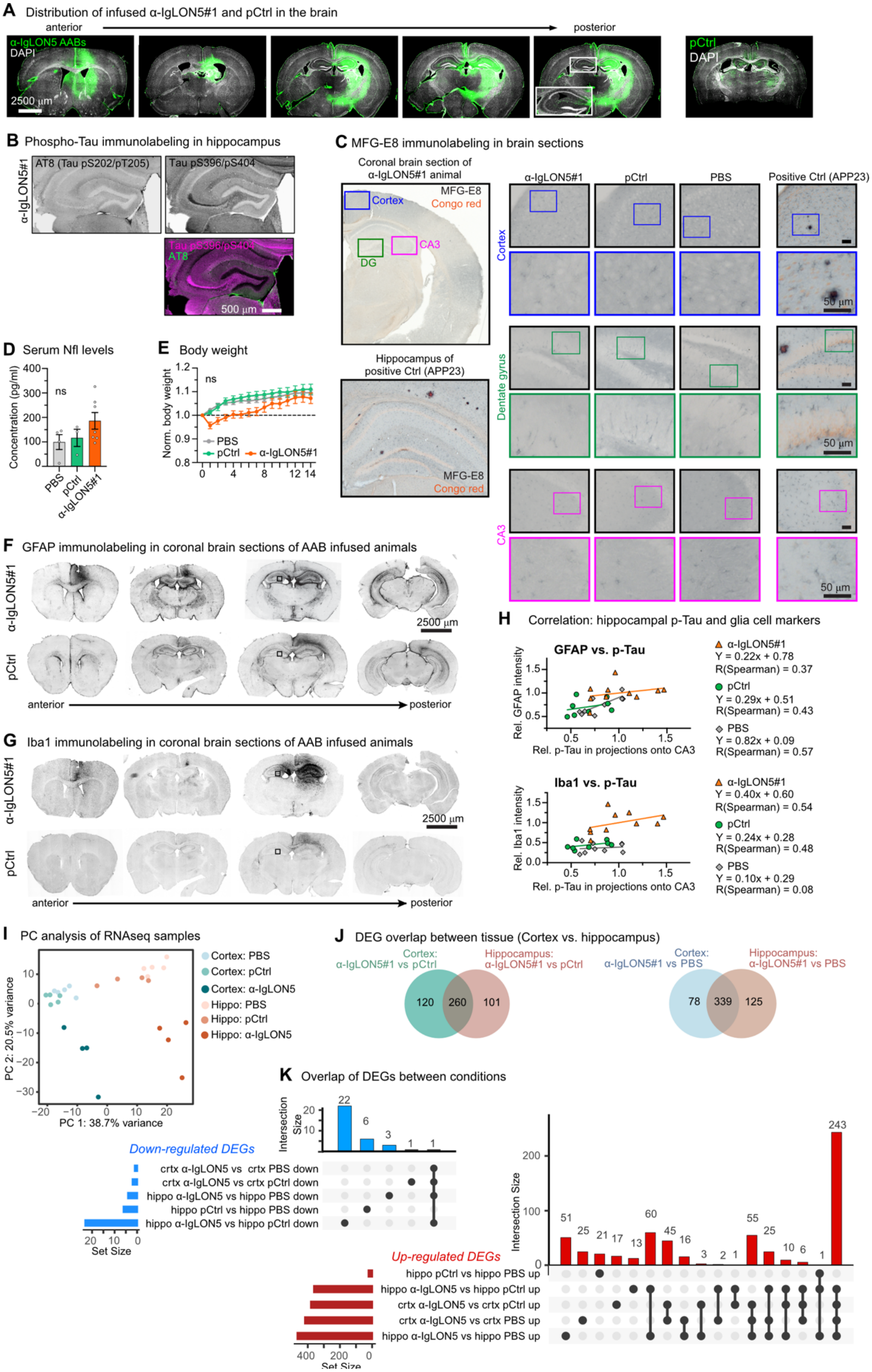
A Distribution of infused α-IgLON5#1 and pCtrl in the brain
anterior
posterior
α-IgLON5 AABs
DAPI
2500 µm
pCtrl
DAPI
B Phospho-Tau immunolabeling in hippocampus
α-IgLON5#1
AT8 (Tau pS202/pT205)
Tau pS396/pS404
Tau pS396/pS404
AT8
500 µm
C MFG-E8 immunolabeling in brain sections
Coronal brain section of α-IgLON5#1 animal
MFG-E8
Congo red
Cortex
CA3
DG
Hippocampus of positive Ctrl (APP23)
MFG-E8
Congo red
α-IgLON5#1
pCtrl
PBS
Positive Ctrl (APP23)
Cortex
Dentate gyrus
CA3
50 µm
50 µm
50 µm
D Serum Nfl levels
Concentration (pg/ml)
400
300
200
100
0
ns
PBS
pCtrl
α-IgLON5#1
E Body weight
Norm. body weight
1.2
1.1
1.0
0.9
0.8
ns
PBS
pCtrl
α-IgLON5#1
0
4
8
12 14
F GFAP immunolabeling in coronal brain sections of AAB infused animals
α-IgLON5#1
pCtrl
2500 µm
anterior
posterior
G Iba1 immunolabeling in coronal brain sections of AAB infused animals
α-IgLON5#1
pCtrl
2500 µm
anterior
posterior
H Correlation: hippocampal p-Tau and glia cell markers
GFAP vs. p-Tau
Rel. GFAP intensity
1.5
1.0
0.5
0.5
1.0
1.5
Rel. p-Tau in projections onto CA3
α-IgLON5#1
Y = 0.22x + 0.78
R(Spearman) = 0.37
pCtrl
Y = 0.29x + 0.51
R(Spearman) = 0.43
PBS
Y = 0.82x + 0.09
R(Spearman) = 0.57
Iba1 vs. p-Tau
Rel. Iba1 intensity
1.5
1.0
0.5
0.0
0.5
1.0
1.5
Rel. p-Tau in projections onto CA3
α-IgLON5#1
Y = 0.40x + 0.60
R(Spearman) = 0.54
pCtrl
Y = 0.24x + 0.28
R(Spearman) = 0.48
PBS
Y = 0.10x + 0.29
R(Spearman) = 0.08
I PC analysis of RNAseq samples
PC 2: 20.5% variance
10
0
−10
−20
−30
−20
−10
0
10
20
PC 1: 38.7% variance
Cortex: PBS
Cortex: pCtrl
Cortex: α-IgLON5
Hippo: PBS
Hippo: pCtrl
Hippo: α-IgLON5
J DEG overlap between tissue (Cortex vs. hippocampus)
Cortex: α-IgLON5#1 vs pCtrl
Hippocampus: α-IgLON5#1 vs pCtrl
120
260
101
Cortex: α-IgLON5#1 vs PBS
Hippocampus: α-IgLON5#1 vs PBS
78
339
125
K Overlap of DEGs between conditions
Intersection Size
20
10
22
6
3
1
1
Down-regulated DEGs
crtx α-IgLON5 vs crtx PBS down
crtx α-IgLON5 vs crtx pCtrl down
hippo α-IgLON5 vs hippo PBS down
hippo pCtrl vs hippo PBS down
hippo α-IgLON5 vs hippo pCtrl down
Set Size
Intersection Size
200
100
0
51
25
21
17
13
60
45
16
3
2
1
55
25
10
6
1
243
Up-regulated DEGs
hippo pCtrl vs hippo PBS up
hippo α-IgLON5 vs hippo pCtrl up
crtx α-IgLON5 vs crtx pCtrl up
crtx α-IgLON5 vs crtx PBS up
hippo α-IgLON5 vs hippo PBS up
400
200
0
Set Size

**Fig. S3. Tau, toxicity, and neuroinflammation in mice infused with AABs.**

(**A**) Representative coronal mouse brain sections (anterior to posterior) showing human antibody distribution (anti-human IgG) upon 14 days of unilateral cerebroventricular α-IgLON5#1 infusion. pCtrl-infused brain section is shown for comparison. (**B**) Representative images showing the hippocampal formation of an α-IgLON5#1 animal immunolabeled for Tau pS202/pT205 (AT8) and Tau pS396/pS404. (**C**) Representative images of cortex, DG and CA3 in α-IgLON5#1 and control brain sections immunolabeled for MFGE-8 and co-stained with Congo red. A brain section of an amyloid-beta expressing APP23 animal is shown as positive control. (**D**) Serum levels of neurofilament light chain (Nfl; by ELISA) are non-significantly elevated in α-IgLON5#1 versus pCtrl and PBS animals. Data shown as mean ± SEM, data points represent individual animals. N = 3 - 7 animals with 3 technical replicates measured per animal. One-way ANOVA with Tukey post-test. (**E**) Body weight relative to Day 0 measured during 14 days of AAB infusion. Data shown as mean ± SEM, N = 8 - 10 animals per group. Student t-test comparing α-IgLON5#1 versus pCtrl or PBS at each time point. (**F**) Representative series of coronal brain sections (anterior to posterior) from α-IgLON5#1 and pCtrl infused animals, immunolabeled for GFAP in astrocytes. Scale bar as indicated. (**G**) Iba1 (microglia) immunolabeling in the same brain sections shown in (F). Scale bar as indicated. (**H**) Correlation (linear regression) between hippocampal GFAP (top) or Iba1 (bottom) fluorescence intensities and Tau pS396/pS404 in projections onto CA3 for α-IgLON5#1, pCtrl, and PBS animals. Data points represent mean of individual animals. (**I**) Principal component analysis (PCA) of RNA-seq data obtained from contralateral cortical and hippocampal tissues of mice infused with PBS (N = 5), pCtrl (N = 4 - 5), or α-IgLON5#1 (N = 4) shows a clear separation of α-IgLON5#1 from control cortices and hippocampi. (**J**) Venn diagrams showing the number of unique and shared differentially expressed genes (DEGs) between cortex and hippocampus for α-IgLON5#1 vs. pCtrl (left), and α-IgLON5#1 vs. PBS (right). (**K**) UpSet plots of intersection analysis of down- (blue) and up- (red) regulated DEGs across experimental groups. The largest overlap (n = 243 DEGs) is found in upregulated DEGs between α-IgLON5#1 cortex and hippocampus vs. respective brain regions in PBS and pCtrl, indicating consistent upregulation of transcripts in hippocampus and cortex of α-IgLON5#1-treated animals. All scale bars as indicated.

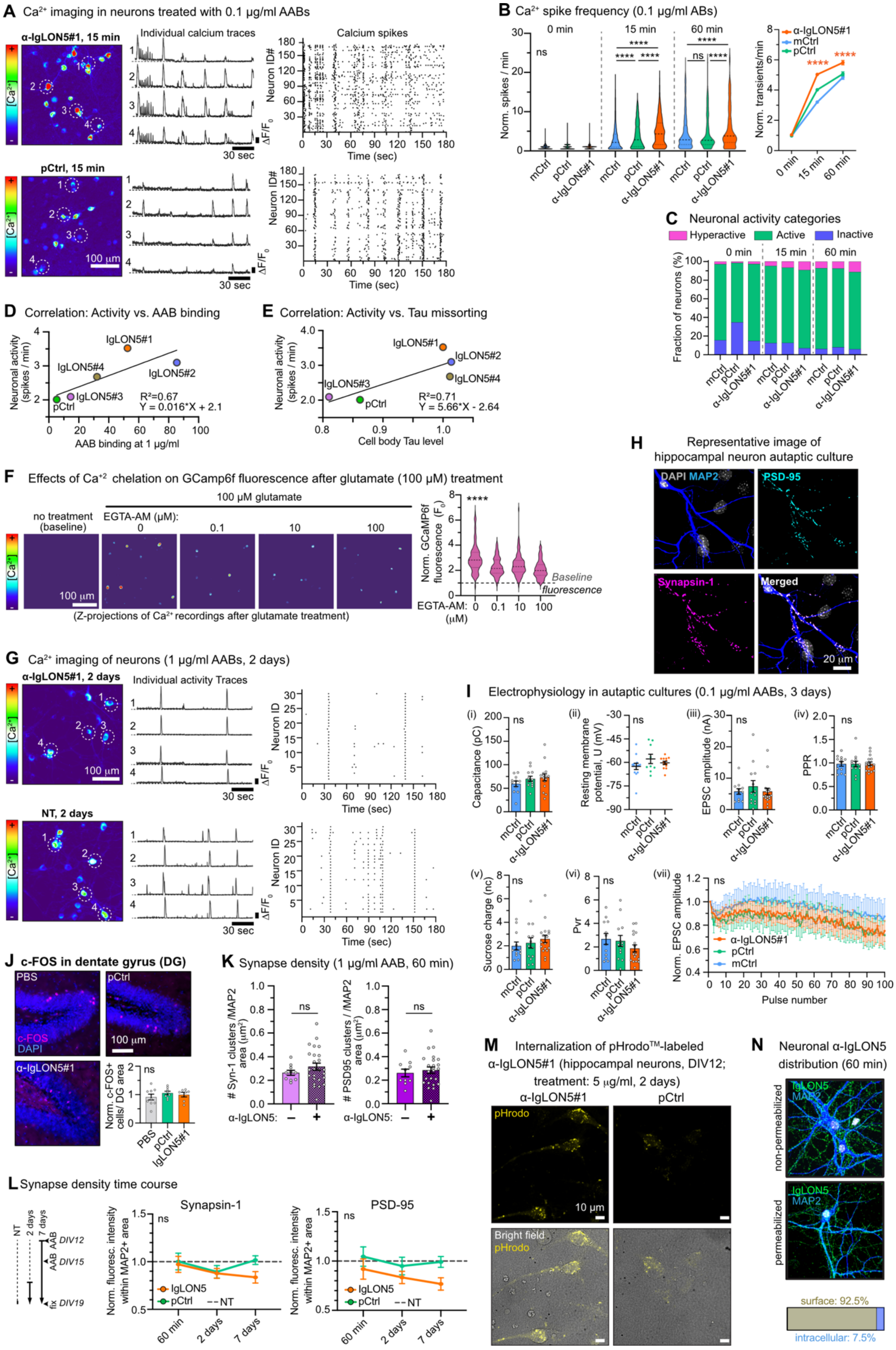

A Ca²⁺ imaging in neurons treated with 0.1 µg/ml AABs
α-IgLON5#1, 15 min
Individual calcium traces
Calcium spikes
Neuron ID#
Time (sec)
30 sec
pCtrl, 15 min
100 µm
B Ca²⁺ spike frequency (0.1 µg/ml ABs)
0 min
15 min
60 min
Norm. spikes / min
ns
****
mCtrl
pCtrl
α-IgLON5#1
Norm. transients/min
C Neuronal activity categories
Hyperactive
Active
Inactive
Fraction of neurons (%)
D Correlation: Activity vs. AAB binding
Neuronal activity (spikes / min)
IgLON5#1
IgLON5#4
IgLON5#2
IgLON5#3
pCtrl
R²=0.67
Y = 0.016*X + 2.1
AAB binding at 1 µg/ml
E Correlation: Activity vs. Tau missorting
R²=0.71
Y = 5.66*X - 2.64
Cell body Tau level
F Effects of Ca⁺² chelation on GCamp6f fluorescence after glutamate (100 µM) treatment
100 µM glutamate
no treatment (baseline)
EGTA-AM (µM):
(Z-projections of Ca²⁺ recordings after glutamate treatment)
Norm. GCaMP6f fluorescence (F0)
Baseline fluorescence
H Representative image of hippocampal neuron autaptic culture
DAPI MAP2
PSD-95
Synapsin-1
Merged
20 µm
G Ca²⁺ imaging of neurons (1 µg/ml AABs, 2 days)
α-IgLON5#1, 2 days
Individual activity Traces
NT, 2 days
Neuron ID
I Electrophysiology in autaptic cultures (0.1 µg/ml AABs, 3 days)
Capacitance (pC)
Resting membrane potential, U (mV)
EPSC amplitude (nA)
PPR
Sucrose charge (nc)
Pvr
Norm. EPSC amplitude
Pulse number
J c-FOS in dentate gyrus (DG)
PBS
pCtrl
c-FOS
DAPI
α-IgLON5#1
Norm. c-FOS+ cells/ DG area
K Synapse density (1 µg/ml AAB, 60 min)
### Syn-1 clusters /MAP2 area (µm²)
### PSD95 clusters / MAP2 area (µm²)
α-IgLON5:
L Synapse density time course
NT
2 days
7 days
AAB
DIV12
DIV15
fix
DIV19
Synapsin-1
PSD-95
Norm. fluoresc. intensity within MAP2+ area
IgLON5
pCtrl
NT
60 min
M Internalization of pHrodo™-labeled α-IgLON5#1 (hippocampal neurons, DIV12; treatment: 5 µg/ml, 2 days)
pHrodo
10 µm
Bright field
N Neuronal α-IgLON5 distribution (60 min)
IgLON5
MAP2
non-permeabilized
permeabilized
surface: 92.5%
intracellular: 7.5%

**Fig. S4. Supplementary Figure S4. Activity of AAB-treated neurons.**

(**A**) Representative images showing $Ca^{2+}$ levels (time-integrated; pseudo-colored) derived from time-lapse recordings of GCamp6f expressing neurons treated with 0.1 μg/ml α-IgLON5#1 or pCtrl for 15 min. White outlined ROIs indicate the position of the neurons for which GCamp6f intensity traces ($\Delta F/F_0$ vs time) are shown. Raster plots show the occurrence of individual $Ca^{+2}$ spikes in cell bodies of all cells in one field of view (N ~ 180 cells, y-axis). Scale bars as indicated. (**B**) Spike rates in neurons treated with 0.1 μg/ml of α-IgLON5#1, pCtrl, or mCtrl (non-reactive monoclonal human antibody mGO (*37*)) for 15 min and 60 min. $Ca^{2+}$ spikes were recorded before (0 min) and after 15 min and 60 min of AAB treatment in the same cultures. Data shown as mean±SEM, N = 3 independent experiments, 71-994 neurons per group. One-way ANOVA with Tukey post-test. (**C**) Percentage of inactive (blue), active (green), and hyperactive (pink) neurons in cultures treated with 0.1 μg/ml α-IgLON5#1, pCtrl and mCtrl for 0 min, 15 min or 60 min. Hyperactive criteria: # of spikes/min > mean+2*SD of pCtrl at each recording time point. Active: 0 < # of spikes/min < mean+2*SD of pCtrl at each recording time point. Silent: no spikes. (**D**) Correlation (linear regression) between neuronal activity and AAB binding after treatment with 1 μg/ml AABs for 60 min. Data points represent α-IgLON5 AABs obtained from different patients. (**E**) Correlation (linear regression) between neuronal activity after treatment with 1 μg/ml AABs for 60 min and cell body Tau levels after treatment with 1 μg/ml AABs for 2 days. Data points represent α-IgLON5 AABs obtained from different patients. (**F**) Representative images showing $Ca^{2+}$ levels (time-integrated; pseudo-colored) derived from time-lapse recordings in GCamp6f -expressing hippocampal neurons, pre-treated with increasing concentrations of EGTA-AM (0.1-100 μM) for 20 min followed by glutamate (100 μM) stimulation for 20 min. Violin plots show GCamp6f fluorescence intensities in neurons. N = 90-124 neurons per group were analyzed. One-way ANOVA with Tukey's post-test. Scale bar as indicated. (**G**) Representative images showing $Ca^{2+}$ levels (time-integrated; pseudo-colored) derived from time-lapse recordings, individual activity traces, and raster plots for detected $Ca^{2+}$ spikes in neurons treated with 1 μg/ml α-IgLON5#1 for 2 days or not treated (NT). Scale bars as indicated. (**H**) Representative image of autaptic cultures immunolabeled for DAPI, MAP2, PSD95 and Synapsin-1. Scale bars as indicated. (**I**) Electrophysiological recordings in autaptic cultures treated with 0.1 μg/ml α-IgLON5 AABs, pCtrl and mCtrl for 3 days. mCtrl is a non-reactive human monoclonal control antibody. Measured parameters are (i) membrane capacitance (pC), (ii) resting membrane potential, (iii) excitatory postsynaptic current (EPSC) amplitude, (iv) EPSC paired-pulse ratio (PPR), (v) readily releasable pool size (sucrose charge), (vi) synaptic vesicle release probability (Pvr), and (vii) EPSC release during 10 Hz stimulation. N = 3 independent autaptic cultures. Data shown as mean±SEM. One-way ANOVA with Tukey post-test. (**J**) Immunolabeling of c-FOS in the DG of α-IgLON5#1 and control animals. Data shown as mean±SEM, N = 7-9 animals, 3-5 brain sections per animal. One-way ANOVA with Tukey post-test. Scale bar = 100 μm. (**K**) Pre- (Synapsin-1) and post- (PSD95) synaptic cluster densities (# punctae/MAP+ area) in neurons treated with 1 μg/ml α-IgLON5#1 for 60 min, or not. N = 3 experiments with 3-5 images analyzed per condition and experiment. Data shown as mean±SEM, Student t-test. (**L**) Analysis of synaptic marker (Synapsin-1 and PSD-95) densities upon incubation of neurons with 1 μg/ml α-IgLON5#1 for 60 min, 2 days, or 7 days. Data shown as mean±SEM, N = 3 experiments with 3 replicates per experiment and 3-5 images per replicate. Dashed lines indicate corresponding synaptic marker levels of untreated neurons at DIV19. (**M**) Representative images showing internalization and lysosomal acidification (= fluorescence) of pHrodoTM-labelled α-IgLON5#1 or pCtrl in primary hippocampal mouse neurons after 2 days treatment. Scale bars indicated. (**N**) Representative images for immunolabeling of cell surface α-IgLON5#1 in unpermeabilized and total α-IgLON5#1 in permeabilized (TX-100) neurons. Percentage of surface vs. internalized α-IgLON5#1 clusters after 60 min of treatment was calculated from 3 images each. Note, image of unpermeabilized neurons is the same as in Fig. 1D**.**

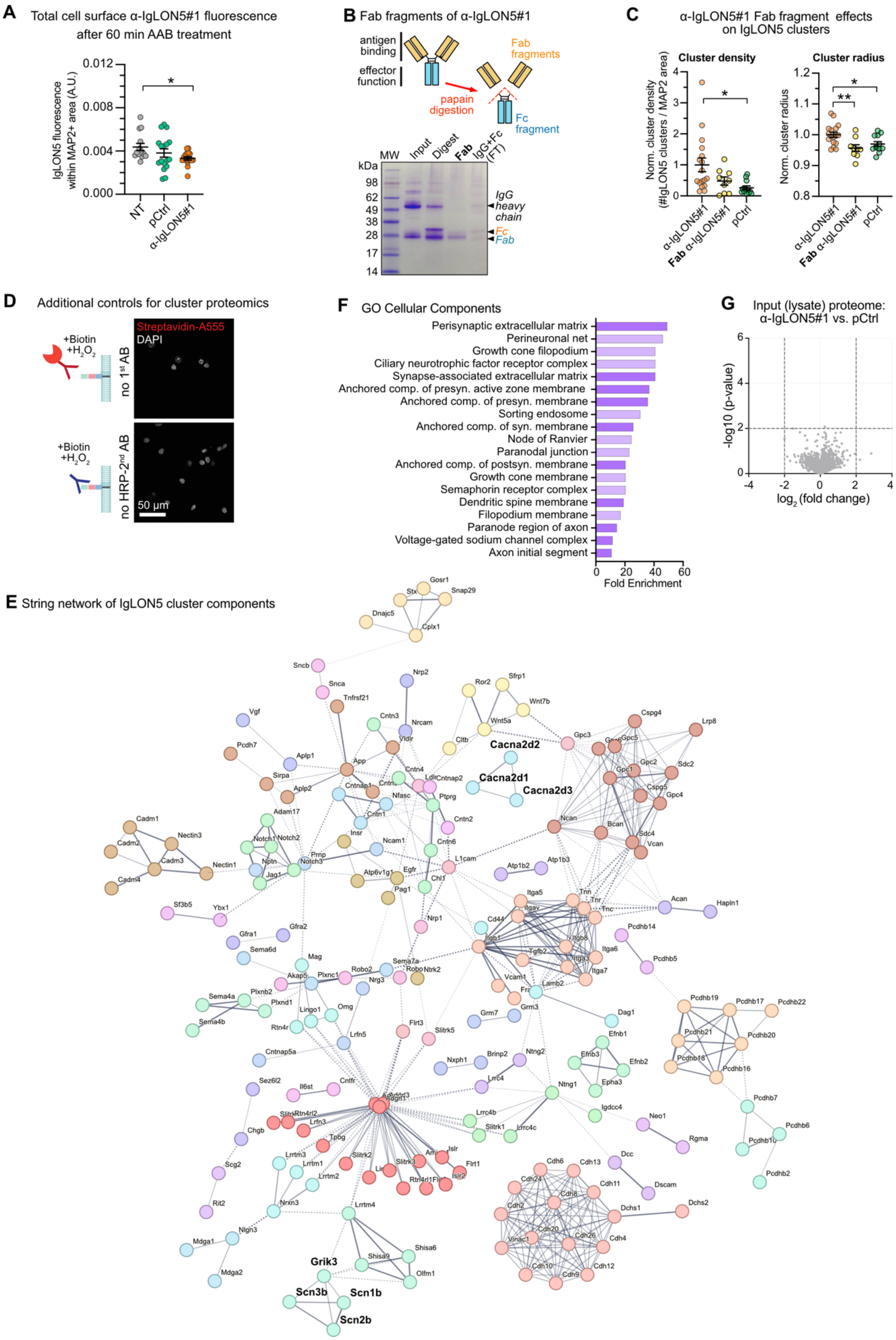
A Total cell surface α-IgLON5#1 fluorescence after 60 min AAB treatment
IgLON5 fluorescence within MAP2+ area (A.U.)
0.012
0.008
0.004
0.000
*
NT
pCtrl
α-IgLON5#1
B Fab fragments of α-IgLON5#1
antigen binding
effector function
Fab fragments
papain digestion
Fc fragment
MW
Input
Digest
Fab
IgG+Fc (FT)
kDa
98
62
49
38
28
17
14
IgG heavy chain
Fc
Fab
C α-IgLON5#1 Fab fragment effects on IgLON5 clusters
Cluster density
Norm. cluster density (#IgLON5 clusters / MAP2 area)
Cluster radius
Norm. cluster radius
α-IgLON5#1
Fab α-IgLON5#1
pCtrl
D Additional controls for cluster proteomics
+Biotin +$H_2O_2$
Streptavidin-A555
DAPI
no 1st AB
no HRP-2nd AB
50 µm
F GO Cellular Components
Perisynaptic extracellular matrix
Perineuronal net
Growth cone filopodium
Ciliary neurotrophic factor receptor complex
Synapse-associated extracellular matrix
Anchored comp. of presyn. active zone membrane
Anchored comp. of presyn. membrane
Sorting endosome
Anchored comp. of syn. membrane
Node of Ranvier
Paranodal junction
Anchored comp. of postsyn. membrane
Growth cone membrane
Semaphorin receptor complex
Dendritic spine membrane
Filopodium membrane
Paranode region of axon
Voltage-gated sodium channel complex
Axon initial segment
0 20 40 60
Fold Enrichment
G Input (lysate) proteome: α-IgLON5#1 vs. pCtrl
-log10 (p-value)
$log_2$ (fold change)
E String network of IgLON5 cluster components
Cacna2d2
Cacna2d1
Cacna2d3
Grik3
Scn3b
Scn1b
Scn2b

**Fig. S5. α-IgLON5 AAB Fab fragments and IgLON5 surface cluster analysis.**

(**A**) Total fluorescence intensity of α-IgLON5#1 on the neuronal cell surface in fixed neurons with and without treatment of IgLON5 prior to fixation. Data shown as mean±SEM, data points represent images analyzed. N = 3 experimental and 3-5 technical replicates. One-Way ANOVA with Tukey post-test. (**B**) Principle and representative reducing SDS-PAGE analysis of Fab fragment generation by papain digestion. Lane-1: input = undigested IgG, Lane-2: papain digested IgG-containing reduced Fc and Fab fragments, Lane-3: purified and reduced IgG Fab fragments (~25 kDa), and Lane-4: flow-through (FT) containing undigested IgG, Fc and Fab fragments. (**C**) Normalized IgLON5 surface cluster density (# IgLON5 clusters/ MAP2+ area) and radius in neurons treated with 1 μg/ml undigested pCtrl or α-IgLON5#1 (~150 kDa, containing 2 Fab fragments per IgG), or with 1.5 μg/ml isolated, non-reduced Fab α-IgLON5#1 (~50 kDa per Fab fragment) for 60 min. N = 2 independent experiments with 3 replicates each and 3 images per replicate. Data shown as mean±SEM. One-way ANOVA with Tukey's post-test. (**D**) Representative images showing no biotinylation in control conditions of IgLON5 surface cluster proximity labeling lacking either the α-IgLON5#1 primary or the HRP-coupled anti-human secondary. (**E**) Visualization of protein-protein interaction network (STRING) of proteins identified in the IgLON5 cluster proteome. (**F**) Gene Ontology (GO) enrichment analysis of cellular components associated with proteins significantly enriched in α-IgLON5 AAB-induced surface clusters. (**G**) Volcano plot for input (total lysate) samples comparing α-IgLON5#1 vs. pCtrl.

**Table S1. Antibody list.**

**IF**: Immunofluorescence    **IHC:** Immunohistochemistry    **WB:** Westernblot

| Primary antibody | Species | Catalog number | company | Application and dilution | | |
|---|---|---|---|---|---|---|
| | | | | IF | IHC | WB |
| MAP2 | chicken | ab92434 | Abcam | 1:1000 | | |
| Synapsin-1 | guinea pig | 106004 | Synaptic Systems | 1:500 | | |
| PSD-95 | mouse | MAB159 | Millipore | 1:1000 | | |
| GFAP | mouse | MAB3402 | Millipore | | 1:500 | |
| Iba1 | rabbit | 019-19741 | WAKO | | 1:500 | |
| CD68 | rat | MCA1957T | BioRad | | 1:500 | |
| Complemet3 (C3) | rat | HM1045 | HycultBiotech | | 1:100 | |
| C1q | rabbit | ab182451 | Abcam | | 1:1000 | |
| CD8 | rat | MA1-145 | Invitrogen | | 1:100 | |
| Caspase-3 | rabbit | 559565 | BD Pharmingen | | 1:500 | |
| MFG-E8 | goat | AF2805 | R&D Systems | | 1:1000 | |
| c-FOS | rat | 226 017 | Synaptic systems | 1:1000 | 1:500 | |
| Biotin | rabbit | 5597S | CellSignaling | | | 1:1000 |
| GAPDH | mouse | CB1001 | Millipore | | | 1:1000 |
| Tau-5 | mouse | MA5-12808 | Invitrogen | 1:1000 | | 1:5000 |
| Tau (DAKO) | rabbit | 10024 | DAKO | | | 1:5000 |
| p-Tau (Ser396)* | rabbit | ab109390 | Abcam | | 1:1000 | 1:5000 |
| pTau (Ser404)* | rabbit | ab92676 | Abcam | | 1:500 | 1:2000 |
| p-Tau (Ser202)** + | rabbit | 39357S | Cell Signaling Technology | | 1:500 | 1:1000 |
| p-Tau (Thr205)** | rabbit | 44-738G | Invitrogen | | 1:500 | 1:1000 |
| p-Tau (Thr231) (AT180) | mouse | MN1040 | Invitrogen | | | 1:500 |
| p-Tau (S202/S205) (AT8) | mouse | MN1020 | Invitrogen | | | 1:1000 |
| p-Tau (Ser199) | rabbit | ab4749 | Abcam | | | 1:1000 |

**Mixed to mimick PHF-1 epitope; **Mixed to mimick AT8 epitope; + This antibody didn't work for IHC*

| Secondary antibody | Catalog number | company | Application and dilution | | |
|---|---|---|---|---|---|
| | | | IF | IHC | WB |
| Goat anti-human 488 | 109-545-003 | Jackson ImmunoResearch | 1:500 | | |
| Goat anti-Mouse, Alexa Fluor 488 | A-21121 | Invitrogen | 1:1000 | | |

| | | | | | |
|---|---|---|---|---|---|
| Goat anti-Rat, Alexa Fluor 488 | A-1106 | Invitrogen | | | |
| Goat anti-Guinea Pig, Alexa Fluor 488 | A-11073 | Invitrogen | | | |
| Goat anti-Rat, Alexa Fluor 555 | A-21434 | Invitrogen | | | |
| Goat anti-Chicken, Alexa Fluor Plus 555 | A-32932 | Invitrogen | | | |
| Goat Anti-Rabbit, Alexa Fluor 647 | A-21245 | Invitrogen | 1:500 | | |
| Streptavidin, Alexa Fluor 555 | S32355 | Invitrogen | 1:500 | | |
| Goat anti-Mouse, Alexa Fluor 790 | A11375 | Invitrogen | | | 1:20 000 |
| Goat anti-Rabbit, Alexa Fluor 790 | A11367 | Invitrogen | | | 1:20 000 |
| Goat anti-Mouse, DyLight 680 | 35519 | Invitrogen | | | 1:10 000 |
| Goat anti-Rabbit, DyLight 680 | 35569 | Invitrogen | | | 1:10 000 |
| Goat anti-human HRP | 31410 | Invitrogen | | 1:200 | |
| | | | 1:1000 (Proximity biotinylation) | | |
